# Gaucher Disease Protects Against Tuberculosis

**DOI:** 10.1101/2022.10.16.512394

**Authors:** Jingwen Fan, Victoria L. Hale, Lindsey T Lelieveld, Laura J. Whitworth, Elisabeth M. Busch-Nentwich, Mark Troll, Paul H. Edelstein, Timothy M. Cox, Francisco J. Roca, Johannes M.F.G. Aerts, Lalita Ramakrishnan

## Abstract

Biallelic mutations in the glucocerebrosidase (*GBA1*) gene cause Gaucher disease, characterized by lysosomal accumulation of glucosylceramide and glucosylsphingosine in macrophages. This and other lysosomal diseases occur with high frequency in Ashkenazi Jews. It has been proposed that the underlying mutations confer a selective advantage, in particular conferring protection against tuberculosis. Here, using a zebrafish Gaucher disease model, we find that the mutation *GBA1* N370S, predominant among Ashkenazi Jews, increases resistance to tuberculosis through the microbicidal activity of glucosylsphingosine in macrophage lysosomes. Consistent with lysosomal accumulation occurring only in homozygotes, heterozygotes remain susceptible to tuberculosis. Thus, our findings reveal a mechanistic basis for protection against tuberculosis by *GBA1* N370S and provide biological plausibility for its selection if the relatively mild deleterious effects in homozygotes were offset by significant protection against tuberculosis, a rampant killer of the young in Europe through the Middle Ages into the 19^th^ century.

**Significance Statement:** Gaucher disease is a recessively inherited disorder in which the lipids glucosylceramide and glucosylsphingosine accumulate in lysosomes of macrophages. Macrophages are the first immune cells to engulf infecting bacteria and we find that glucosylsphingosine increases their ability to kill *Mycobacterium tuberculosis* that causes tuberculosis. Gaucher disease due to a particular mutation is frequent in Ashkenazi Jews. Since from the middle ages they were often confined to areas of high tuberculosis prevalence, it has been proposed that the mutation prevailed because heterozygotes, who do not accumulate lipids nor manifest Gaucher disease, were protected. Our findings raise the possibility that selection operated on homozygotes manifesting mild forms of Gaucher disease who were protected against tuberculosis which would often have been fatal.

## Introduction

Tuberculosis (TB) features multiple interactions of *Mycobacterium tuberculosis* (Mtb) with host macrophages, each with the potential to determine if the infection will progress or be cleared (1). Zebrafish develop TB-like disease when infected with their natural pathogen *Mycobacterium marinum* (Mm), a close relative of Mtb (1). In particular, the optically transparent, and genetically and pharmacologically tractable zebrafish larva has enabled delineation of the early steps of TB pathogenesis and the host-mycobacterium interactions that shape them, with zebrafish findings providing insights into human TB pathogenesis and treatment and forming the basis for pre-clinical and clinical studies and clinical trials (1–14).

Through a zebrafish genetic screen, we previously identified a mutant with a lysosomal storage disorder due to a deficiency in lysosomal cysteine cathepsins that was hypersusceptible to mycobacterial infection (7). We found that lysosomal storage in macrophages causes hypersusceptibility by impairing their migration into the developing tuberculous granuloma, causing pathological necrosis which promotes mycobacterial growth (7). In contrast to the ultra-rare cathepsin deficiencies causing accumulation of proteinaceous material, lysosomal diseases that impair recycling of lipids are far more prevalent (15). Accordingly, we sought to determine whether a lysosomal disease associated with pathological storage of lipids affected susceptibility to TB. Glucocerebrosidase (GBA) deficiency, which causes Gaucher disease, is one of the most common lysosomal disorders (15, 16), and is of particular interest because it principally affects macrophages, host cells that interact early and critically with mycobacteria. In Gaucher disease, the lysosomal compartment of macrophages expands and becomes engorged with sphingolipid to assume a storage phenotype (17, 18). Moreover, these cells, widely known as Gaucher cells, have been shown to have defective migration (17, 18).

The zebrafish was an ideal model in which to address the question of how the macrophage lysosomal storage of Gaucher disease might impact TB; over the last few years, it has come into its own as a facile model for Gaucher disease that recapitulates its sphingolipid accumulation and key multisystem pathologic manifestations – hematopoietic, including the hallmark macrophage lysosomal storage, visceral, bony and skeletal and neuronopathic (19–23). The use of activity-based probes and mass spectrometric techniques have enabled the detailed biochemical assays to confirm in the zebrafish Gaucher disease model the metabolic shifts seen in human Gaucher disease (21). Importantly, the use of the zebrafish model has enabled the clarification of the role of the two glucocerebrosidases GBA1 (lysosomal facing) and GBA2 (cytosolic facing) and the downstream enzyme lysosomal acid ceramidase in lipid accumulation and disease phenotypes (21, 24).

Based on our findings with the cathepsin mutant, our prediction was that GBA-deficient zebrafish would be hypersusceptible to mycobacterial infection. Thus we were surprised to find that GBA-deficient zebrafish larvae were resistant to both Mm and Mtb, despite the fish having cardinal manifestations of human Gaucher disease, particularly overt macrophage lysosomal storage and accompanying migration defects. We have delineated the resistance mechanism and shown it to be operant from very early in infection and relevant in the context of the common Ashkenazi Jewish N370S Gaucher disease allele (25–28). Our findings shed light on the decades-long debate about the persistence of this allele – selection versus founder effect resulting in genetic drift (29–33). They provide biological evidence in favor of its TB-driven selection over the centuries.

## Results

### *gba1* mutant zebrafish develop human Gaucher disease manifestations

We examined a zebrafish mutant in the orthologous gene *gba1* (*gba1^sa1621^*) with a premature stop mutation in the region encoding the GBA catalytic domain (*SI Appendix*, Fig. S1*A* and S1*B*). Like the cathepsin-deficient zebrafish (7), at 3 days post-fertilization (dpf), *gba1^sa1621/sa1621^*mutants had an increased proportion of enlarged, rounded brain-resident macrophages (microglia) with Lysotracker staining showing enlarged lysosomes containing accumulated cell debris as evidenced by acridine orange staining (Fig. 1A-1D). This macrophage phenotype is similar to that described for human Gaucher disease, and like human *GBA1* heterozygotes, *gba1^sa1621^*heterozygotes had normal macrophages (17, 18) (Fig. 1*A*-1*D*). Macrophages manifesting lysosomal storage moved slowly, as expected (7) (Fig. 1*E*). *gba1^sa1621^* homozygotes grew into early adulthood but were smaller in size and had curved spines that were previously reported for other zebrafish *gba1* mutants, recapitulating the growth retardation and kyphosis seen in many Gaucher disease patients (19, 20, 24, 34) (Fig. 1*F*). Moreover, between 75 and 80 days of age, all exhibited abnormal swimming characterized by a spinning motion (Supplementary Movie S1) (19). This indicates involvement of the nervous system, which also complicates severe human Gaucher disease (34). None of the wild-type or heterozygous siblings manifested any of these pathological phenotypes during the 2 to 2.5-year observation period.

**Fig. 1.**
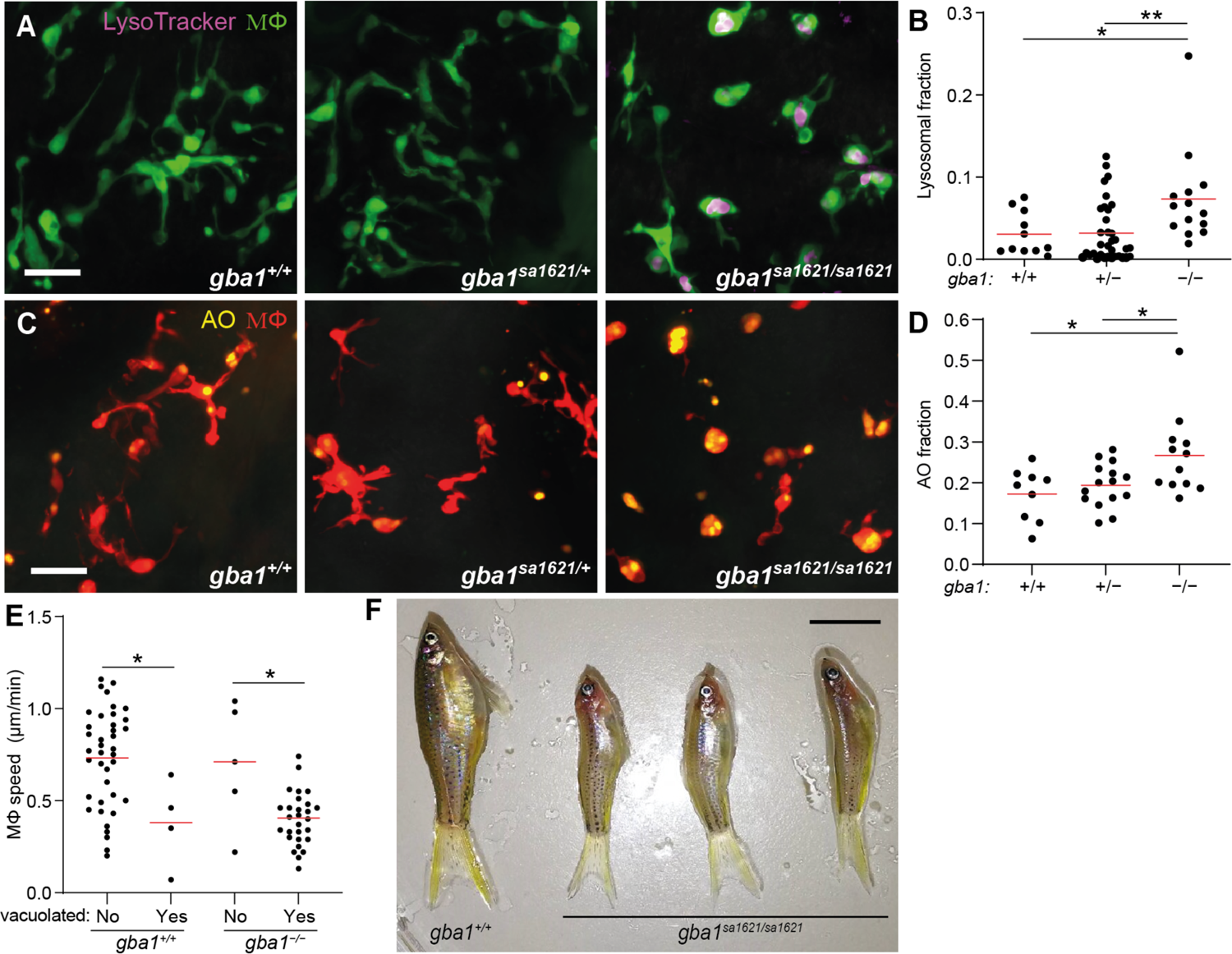
*gba1* mutant zebrafish manifests Gaucher disease. *(A)* Maximum intensity projection of pseudo-colored representative confocal images of YFP-expressing macrophages stained with LysoTracker Red in 3 dpf zebrafish brains. Scale bar, 20 μm. *(B)* LysoTracker Red volume per macrophage in 3 dpf brains. Each point represents the mean volume fraction per macrophage in each brain. Horizontal red bars, means. *\*P <* 0.05, \**\*P* < 0.01 (one-way ANOVA with Tukey’s post-test). Representative of at least 3 independent experiments. *(C)* Maximum intensity projection of representative confocal images of tdTomato-expressing macrophages stained with acridine orange (AO) in 3 dpf zebrafish brains. Scale bar, 20 μm. *(D)* AO volume per macrophage in the brains of 3 dpf zebrafish. Each point represents the average AO volume fraction per macrophage in each fish. Horizontal red bars, means. *\*P <* 0.05 (one-way ANOVA with Tukey’s post-test). *(E)* Homeostatic migration speed of normal and vacuolated macrophages in the brains of 3 dpf zebrafish. Each point represents the mean speed of individual macrophage from the same animal per indicated genotype within 2 hours of observation. Horizontal red bars, means. *\*P <* 0.05 (one-way ANOVA with Tukey’s post-test). Representative of 2-3 animals for each genotype. *(F)* Representative images of three *gba1^sa1621/sa1621^*fish and their wild-type sibling at 77 days post-fertilization (dpf). Scale bar, 1 cm.

### *gba1* mutant zebrafish are resistant to Mm infection

We next tested the susceptibility of *gba1* mutants to mycobacteria. We had previously found that an antisense *gba1* morpholino produced macrophage lysosomal storage and increased susceptibility, similar to the cathepsin-deficient mutants (7) However, because increased susceptibility can be a nonspecific effect from morpholino toxicity and/or off-target effects (35), we sought to confirm these results using genetic mutants. We infected 2 dpf animals from a *gba1^sa1621^*heterozygote incross with fluorescent Mm in the hindbrain ventricle (HBV), an epithelium-lined cavity where mycobacteria interact initially with first-responding resident macrophages before monocyte-mycobacterium interactions become dominant as granulomas form (36) (Fig. 2*A*). Contrary to expectation, we found that they were Mm resistant. Fewer bacteria were present in the HBV at 3 days post-infection (dpi) than in their wild-type siblings (Fig. 2*B* and 2*C*). Heterozygotes had wild-type bacterial burdens (Fig. 2*C*). Intravenous injection of bacteria into the caudal vein (CV) where mycobacteria interact directly with monocytes (36) (Fig. 2*A*) also showed the same resistance phenotype (Fig. 2*D* and 2*E*). To ensure that the resistance phenotype was specifically due to the *gba1* mutation, we created *gba1* G0 crispants using three different guide RNAs (*SI Appendix*, Fig. S1*A*). The pooled *gba1* G0 crispants were also resistant to Mm infection (Fig. 2*F*). We then generated two individual mutants *gba1^cu41^* and *gba1^cu42^* (*SI Appendix*, Fig. S1*A* and S1*B*). Like the *gba1^sa1621^* mutants, both mutants were small with curved spines and developed abnormal swimming around 80 days. Likewise, *gba1^cu41^* and *gba1^cu42^* homozygotes were resistant to Mm infection (Fig. 2*G* and 2*H*). Compound *sa1621*/*cu42* heterozygotes were also resistant (Fig. 2*I*) and restoration of GBA using zebrafish *gba1* RNA eliminated resistance of *gba1* mutants (Fig. 2*J*), confirming that the *gba1* mutation caused resistance.

**Fig. 2.**
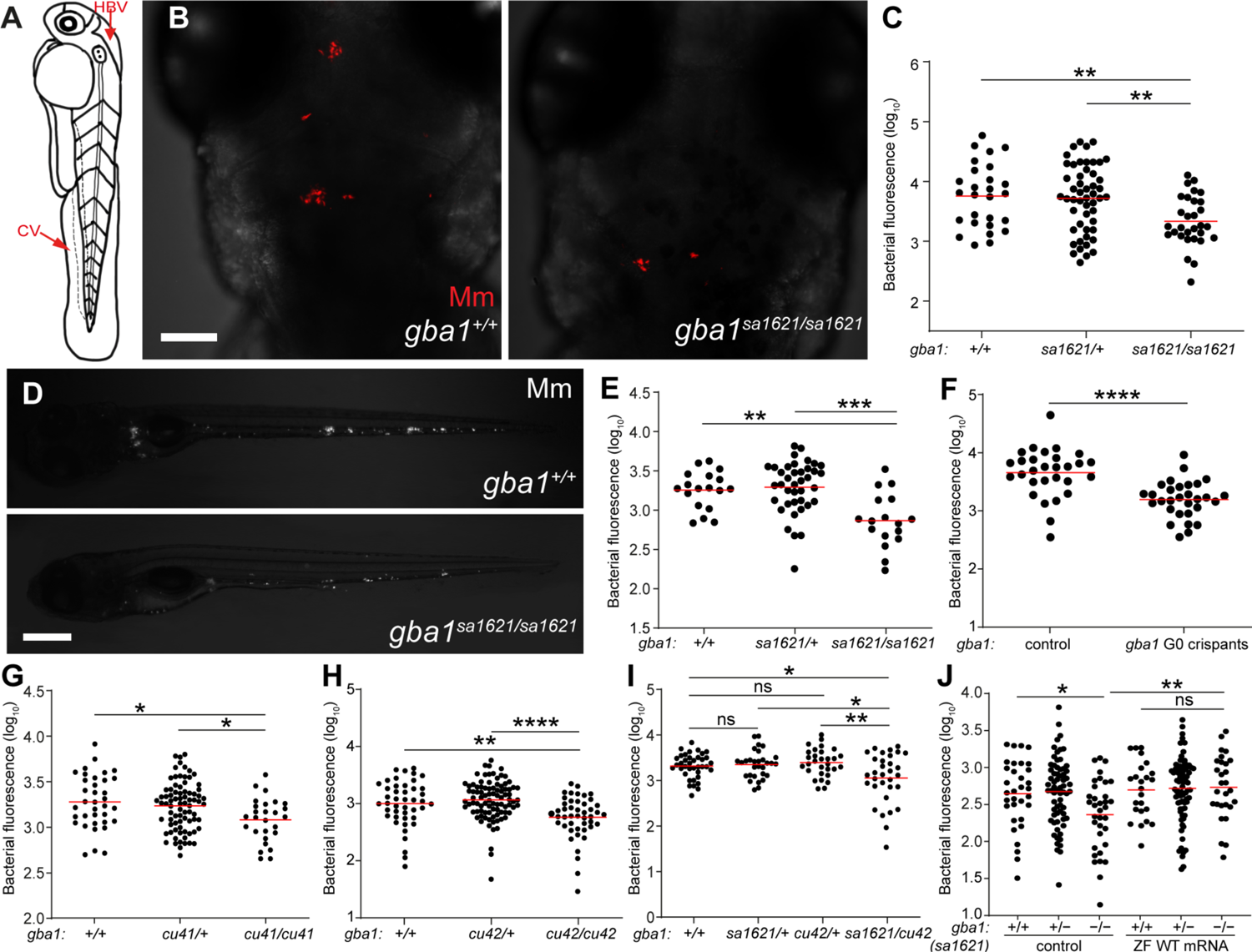
*gba1* mutant zebrafish are resistant to Mm infection. *(A)* Illustration of a zebrafish larva showing the hindbrain ventricle (HBV) and caudal vein (CV) injection sites. *(B)* Maximum intensity projection of representative confocal images of zebrafish larval HBV at 3 dpi after HBV infection with 100-150 Mm. Scale bar, 80 μm. *(C)* Quantification of HBV bacterial burden measured by fluorescence per animal from *(B)*. Horizontal red bars, means; \**\*P <* 0.01 (one-way ANOVA with Tukey’s post-test). Representative of more than 3 independent experiments. *(D)* Representative images of zebrafish larvae at 5dpi after CV infection with 200-300 Mm. Scale bar, 300 μm. *(E)* Quantification of the bacterial burden measured by fluorescence per animal from *(D)*. Horizontal bars, means; \**\*P <* 0.01; \*\**\*P <* 0.001 (one-way ANOVA with Tukey’s post-test). Representative of more than 3 independent experiments. *(F)* Bacterial burden measured by fluorescence per animal in wild-type and *gba1* G0 crispants at 5 dpi after CV infection with 200-300 Mm. Horizontal bars, means; \*\*\**\*P <* 0.0001 (Student’s unpaired t-test). Representative of more than 3 independent experiments. *(G)* Bacterial burden measured by fluorescence in 5 dpi *gba1^cu41/+^*incross larvae after CV infection with 200-300 Mm. Horizontal bars, means; *\*P <* 0.05 (one-way ANOVA with Tukey’s post-test). Representative of 2-3 independent experiments. *(H)* Bacterial burden measured by fluorescence in 5 dpi *gba1^cu42/+^* incross larvae after CV infection with 200-300 Mm. Horizontal bars, means; \**\*P <* 0.01; \*\*\**\*P <* 0.0001 (one-way ANOVA with Tukey’s post-test). Representative of more than 3 independent experiments. *(I)* Bacterial burden measured by fluorescence in wild-type, *sa1621* and *cu42* heterozygotes and *sa1621/cu42* compound mutant siblings at 5 dpi after CV infection with 200-300 Mm. Horizontal bars, means; ns, not significant; \**P <* 0.05; \**\*P <* 0.01 (one-way ANOVA with Tukey’s post-test). *(J)* Bacterial burden measured by fluorescence in *gba1^sa1621/+^* incross larvae, injected with zebrafish WT *gba1* mRNA (200 ng/μl) or vehicle control, 5 dpi after CV infection with 200-300 Mm. Horizontal bars, means; ns, not significant; *\*P <* 0.05; \**\*P <* 0.01 (one-way ANOVA with Tukey’s post-test). Representative of 2 independent experiments.

### GBA-deficient macrophages have increased mycobactericidal capacity

How could the migratory defect of *gba1* mutant macrophages be reconciled with resistance rather than susceptibility to Mm? We reasoned that resistance must manifest at the earliest stage of infection before the shortfall of recruited macrophages that leads to granuloma breakdown from macrophage necrosis. If so, then GBA-deficient macrophages must be better able to restrict growth of the infecting mycobacteria, enabling rapid reduction of the infection burden even before the granuloma stage. To test if this was the case, we used velaglucerase alfa, a mannose-terminated human GBA protein that is taken up into macrophages and an effective enzyme replacement therapy for patients with Gaucher disease (37). Administration of velaglucerase alfa eliminated mutant resistance suggesting that GBA-deficient macrophages have increased intrinsic ability to restrict mycobacterial growth (Fig. 3*A*).

**Fig. 3.**
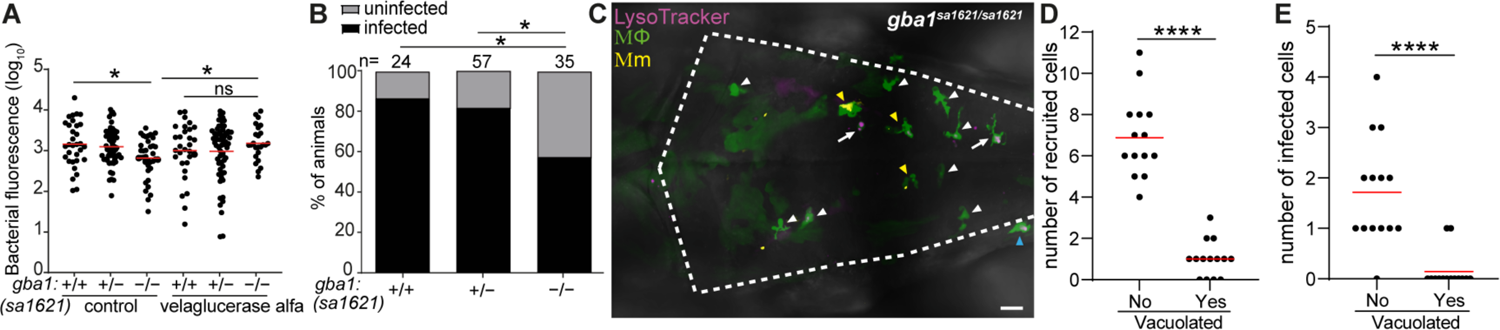
*gba1* mutant macrophages are bactericidal. *(A)* Bacterial burden measured by fluorescence in 3dpi *gba1^sa1621/+^*incross larvae treated with 100 units/ml velaglucerase alfa or vehicle control after HBV infection with 100-150 Mm. Horizontal bars, means; ns, not significant; *\*P <* 0.05 (one-way ANOVA with Tukey’s post-test). Representative of 2 independent experiments. *(B)* Percentage of infected larvae at 5 dpi after HBV infection with a single Mm bacterium. *\*P <* 0.05 (Fisher’s exact test). Representative of 2 independent experiments. *(C)* Maximum intensity projection of pseudo-colored representative confocal image of YFP-expressing macrophages stained with LysoTracker Red in the brain of 2 dpf *gba1* mutant larvae at 6 hours post infection (hpi) in HBV. Arrow marks the vacuolated macrophages. White arrowhead marks non-vacuolated macrophages without phagocytosed Mm. Yellow arrowhead marks non-vacuolated macrophages with phagocytosed Mm. Blue arrowhead marks macrophage that is not recruited in HBV. The majority of the not recruited macrophages are in midbrain. Scale bar, 20 μm. *(D)* Quantification of non-vacuolated and vacuolated macrophages that are recruited to HBV in *gba1* mutant larvae from *(C)*. Each point represents the total number of recruited non-vacuolated or vacuolated macrophages in each fish. Horizontal red bars, means. \*\*\*\**P* < 0.0001 (Student’s unpaired t-test). *(E)* Quantification of non-vacuolated and vacuolated macrophages with phagocytosed Mm from *(C)*. Each point represents the total number of infected non-vacuolated or vacuolated macrophages in each fish. Horizontal red bars, means. \*\*\*\**P* < 0.0001 (Student’s unpaired t-test).

To determine if this reflected an increased ability to kill mycobacteria, i.e., increased microbicidal capacity, we performed an “infectivity assay” where we infected zebrafish in the HBV with a single Mm (36). Because mycobacteria are rapidly phagocytosed by macrophages, the frequency of animals with infection progression versus clearance at 5 dpi is a reliable indicator of macrophage microbicidal capacity (36) (*SI Appendix*, Fig. S2). Significantly more *gba1* mutant zebrafish had cleared infection, confirming that their macrophages were more microbicidal to mycobacteria (Fig. 3*B*). Thus, GBA-deficient brain-resident macrophages possess increased microbicidal capacity that causes increased killing of phagocytosed Mm.

Since we had observed the resistance phenotype after CV infection where the mycobacteria are phagocytosed by blood monocytes of the caudal hematopoietic tissue (CHT) (Fig. 2*D* and 2*E*), these blood monocytes must also be more microbicidal. However, the CHT monocytes did not manifest overt lysosomal storage (*SI Appendix*, Fig. S3). (This difference is likely because the brain macrophages are constantly engulfing apoptotic neurons during brain development at this stage (38)). This finding suggested that GBA-deficient macrophages and monocytes have increased microbicidal capacity even if they have not developed the overt storage phenotype of the pathological Gaucher cell (17, 18).

This model would be consistent with our prior findings that macrophages with advanced lysosomal storage cannot migrate to mycobacteria and participate in the infection (7). We had already shown that in the *gba1* mutant, the vacuolated macrophages with obvious lysosomal storage had a homeostatic migration defect (Fig. 1*E*). We now tested if they also had defective migration to infecting mycobacteria. We infected Mm into the HBV, which first recruits proximate brain-resident macrophages (36). Whereas the majority of macrophages in *gba1* mutant brains had the Gaucher cell phenotype (Fig. 1*A*), these were a distinct minority among those that were recruited to infection (Fig. 3*C* and 3*D*). Rather, infection almost exclusively recruited the few macrophages that were phenotypically normal (Fig. 3*C* and 3*D*). Consequently, these cells were the ones most likely to be infected (Fig. 3*C* and 3*E*). Thus, in GBA-deficient animals, both tissue-resident macrophages and monocytes have enhanced mycobactericidal capacity even if they do not manifest an obvious lysosomal storage phenotype.

### Glucosylsphingosine, which accumulates in GBA-deficient macrophage lysosomes, is mycobactericidal in vitro

One explanation for the increased mycobactericidal capacity of GBA-deficient macrophages was that their accumulated lysosomal products could be directly microbicidal to mycobacteria. We first tested whether the accumulated lysosomal products could account for the increased microbicidal capacity of GBA-deficient macrophages. GBA deficiency causes the accumulation of glucosylceramide which is converted to glucosylsphingosine by lysosomal acid ceramidase (39–41) (Fig. 4*A*).

**Fig. 4.**
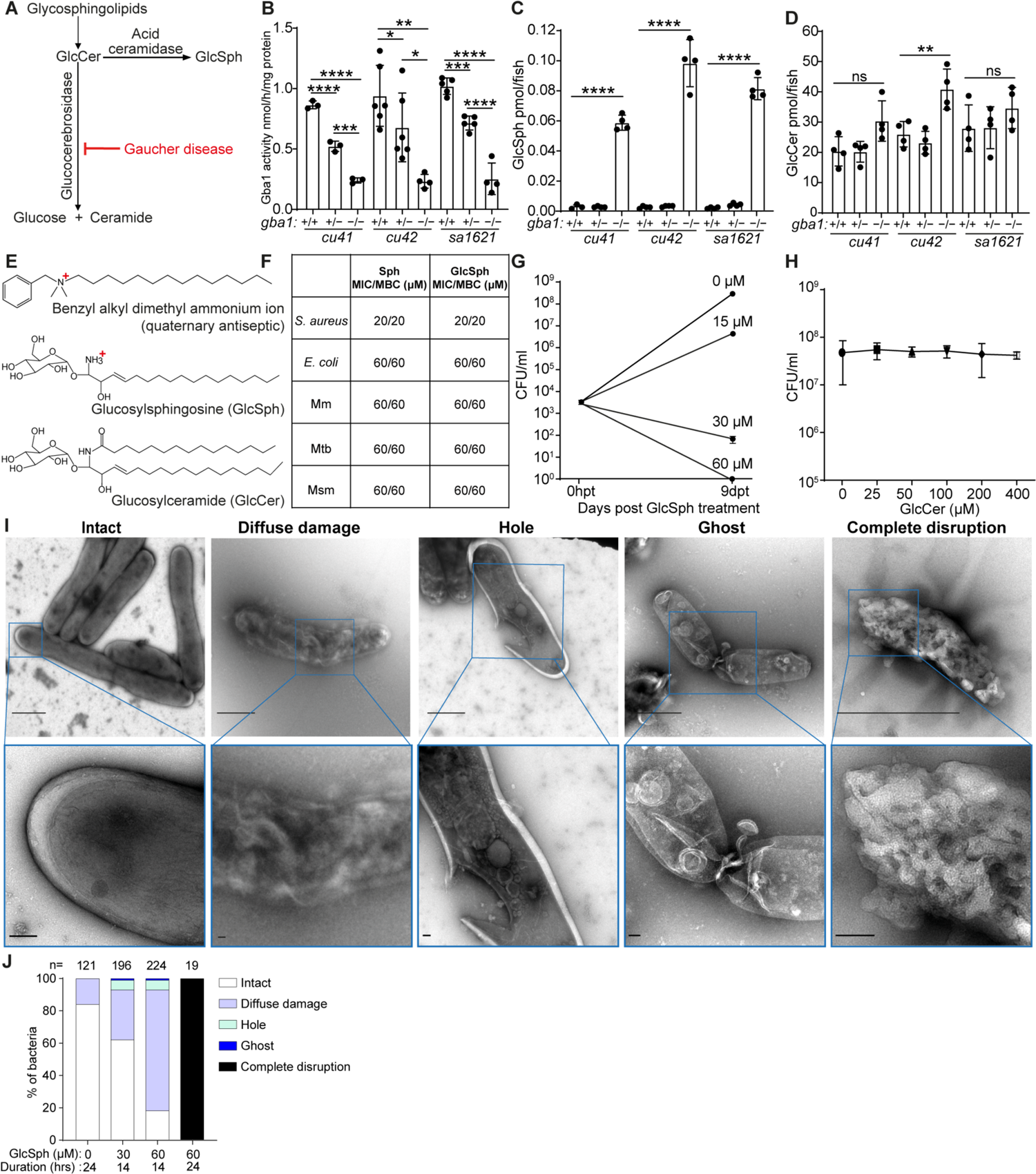
Glucosylsphingosine is bactericidal in vitro. *(A)* Representation of the pathway leading to glucosylceramide and glucosylsphingosine accumulation in Gaucher disease. GlcCer, glucosylceramide. GlcSph, glucosylsphingosine. *(B)* Glucocerebrosidase enzyme activity (nmol/mg) in 5 dpf *gba1 cu41, cu42* and *sa1621* mutants and their wild-type siblings. Mean +/- S.D; *\*P <* 0.05; \**\*P <* 0.01; \*\**\*P <* 0.001; \*\*\**\*P <* 0.0001 (one-way ANOVA with Tukey’s post-test). *(C)* GlcSph (pmol/fish) in 5 dpf *gba1 cu41, cu42* and *sa1621* mutants and their wild-type siblings. Mean +/- S.D; \*\*\**\*P <* 0.0001 (one-way ANOVA with Tukey’s post-test). Representative of 2 independent experiments. *(D)* GlcCer (pmol/fish) in 5 dpf *gba1 cu41, cu42* and *sa1621* mutants and their wild-type siblings. Mean +/- S.D; ns, not significant; \**\*P <* 0.01 (one-way ANOVA with Tukey’s post-test). Representative of 2 independent experiments. *(E)* Chemical structures of benzyl alkyl dimethyl ammonium ion, GlcSph and GlcCer. *(F)* MIC/MBC table of *S. aureus*, *E. coli,* Mm, Mtb and *M. smegmatis* treated with Sph and GlcSph. Minimum inhibitory concentration (MIC) and minimum bactericidal concentration (MBC) determinations were performed according to the Clinical and Laboratory Standards Institute (CLSI) instructions, with modifications for mycobacteria, as described in the materials and methods section. Representative of 1-2 independent experiments performed in duplicate. *(G)* Mm killing by GlcSph. Mean CFU/ml; Vertical bars, upper and lower values of the two technical replicates. *(H)* Mean Mm (CFU/ml) after incubation with increasing GlcCer concentrations for 9 days. Vertical bars, upper and lower values of the two technical replicates. (Starting concentration 2.8×10^3^ CFU/ml). Representative of 2 independent experiments. *(I)* Negative stain-transmission electron microscopy (TEM) images of Msm treated with GlcSph, representative of the types of damage seen. Scale bar, 1 μm (top panels), 100nm (bottom panels). *(J)* Quantification of the types of damage seen in (H) for various GlcSph concentrations.

Glucosylsphingosine is consistently elevated in Gaucher disease macrophage lysosomes and its increased concentration in blood is used as a diagnostic biomarker (39, 42). We confirmed that at 5 dpf all three zebrafish mutants had reduced GBA activity and increased concentrations of glucosylsphingosine, as expected from a similar analysis of another zebrafish *gba1* null mutant (21) (Fig. 4*B* and 4*C*). Also, as in the previously published mutant, neither glucosylceramide nor other sphingolipid concentrations were invariably elevated in these mutants (21) (Fig. 4*D* and *SI Appendix*, Fig. S4). The partial reduction of enzymatic activity in the heterozygotes did not result in any glucosylsphingosine accumulation, just as in the published zebrafish mutant and in human carriers (21, 39) (Fig. 4*B* and 4*C*).

Free sphingosines are broadly microbicidal to Gram-positive and Gram-negative bacteria in vitro and are thought to contribute to the antimicrobial activity of normal skin (43). This is expected as they are cationic surfactants, similar to cationic antiseptics such as the quartenary ammonium compounds, long known to be bactericidal, as the positively charged groups are strongly attracted to negatively charged groups in bacterial membranes, where they impair fluidity, cause leakage, and ultimately disrupt the membranes (44, 45) (Fig. 4*E*). Consistent with its predicted bactericidal activity, sphingosine had the same minimum inhibitory concentration (MIC) and minimum bactericidal activity (MBC) against the Gram-positive bacterium *Staphylococcus aureus* and the Gram-negative bacterium *Escherichia coli* (46) (Fig. 4*F*). Consistent with the membrane-disrupting activity of cationic surfactants, sphingosine was broadly bactericidal to all three mycobacterial species tested, Mm, Mtb and *Mycobacterium smegmatis* (Msm), a rapidly growing environmental nonpathogenic species (Fig. 4*F*). The lower glucosylsphingosine MIC for *S. aureus* (20 μM) than for *E. coli* and mycobacteria (60 μM) may be because the latter bacteria have a multilayer membrane, which could increase resistance to its membrane-disrupting action (47). Importantly, glucosylsphingosine had the same bactericidal activity as sphingosine to all the bacteria tested (Fig. 4*F*). At concentrations lower than its MBC, glucosylsphingosine was still growth inhibitory to Mm, slightly inhibiting and completely inhibiting growth at 15 and 30 μM concentrations, respectively (Fig. 4*G*). Moreover, as predicted because it is a neutral lipid, glucosylceramide did not inhibit bacterial growth even at the highest concentration tested, 400 μM (Fig. 4*E* and 4*H*).

Consistent with its membrane-disrupting action, sphingosine has been shown to produce disruption of the *S. aureus* cell wall (43). We confirmed by negative stain transmission electron microscopy that glucosylsphingosine likewise caused ultrastructural damage to *S. aureus* membranes (*SI Appendix*, Fig. S5). Using Msm (a containment level 1 mycobacterium suitable for use in a shared electron microscopy facility, we then showed that glucosylsphingosine caused similar damage to mycobacterial membranes in a concentration- and time-dependent manner (Fig. 4*I* and 4*J*). Thus, glucosylsphingosine, but not glucosylceramide, is bactericidal to mycobacteria in vitro through its membrane-disrupting activity. This finding is consistent with the membrane-disrupting mycobactericidal activity of glucosyl sphingosine being responsible for the increased macrophage microbicidal capacity of GBA-deficient macrophages we had observed in vivo.

### Accumulated lysosomal glucosylsphingosine is responsible for enhanced mycobacterial killing by GBA-deficient macrophages in vivo

Alternatively, or additionally, resistance in vivo could be an indirect consequence of GBA deficiency, attributable to induction of microbicidal lysosomal enzymes (e.g., lysozyme) and cytokines (e.g., tumor necrosis factor) characteristic of Gaucher disease (1, 15, 48). Therefore, we performed experiments to determine if the accumulated glucosylsphingosine was necessary and sufficient for the increased macrophage microbicidal capacity of GBA-deficient macrophages in vivo. If so, then resistance of *gba1* mutants should be abolished by blocking conversion of the accumulating glucosylceramide to glucosylsphingosine by inhibition of acid ceramidase (Fig. 4*A*). Pharmacological inhibition of acid ceramidase with carmofur (49), eliminated *gba1* mutant resistance (Fig. 5*A*). Next, we superimposed a genetic mutation in acid ceramidase (zebrafish *asah1b*) on *gba1* mutants by creating *gba1* G0 crispants in a cross of *asah1b* heterozygotes. The *gba1-asahb1* double mutants did not accumulate glucosylsphingosine, as shown before (Fig. 5*B*) (24). Importantly, they lost resistance to Mm (Fig. 5*C*). The *asah1b* heterozygotes had residual accumulation of glucosylsphingosine and this was sufficient to confer resistance (Fig. 5*B* and 5*C*). Thus, the increased resistance of *gba1* mutant animals is specifically due to the accumulated glucosylsphingosine, which is necessary for resistance in vivo.

**Fig. 5.**
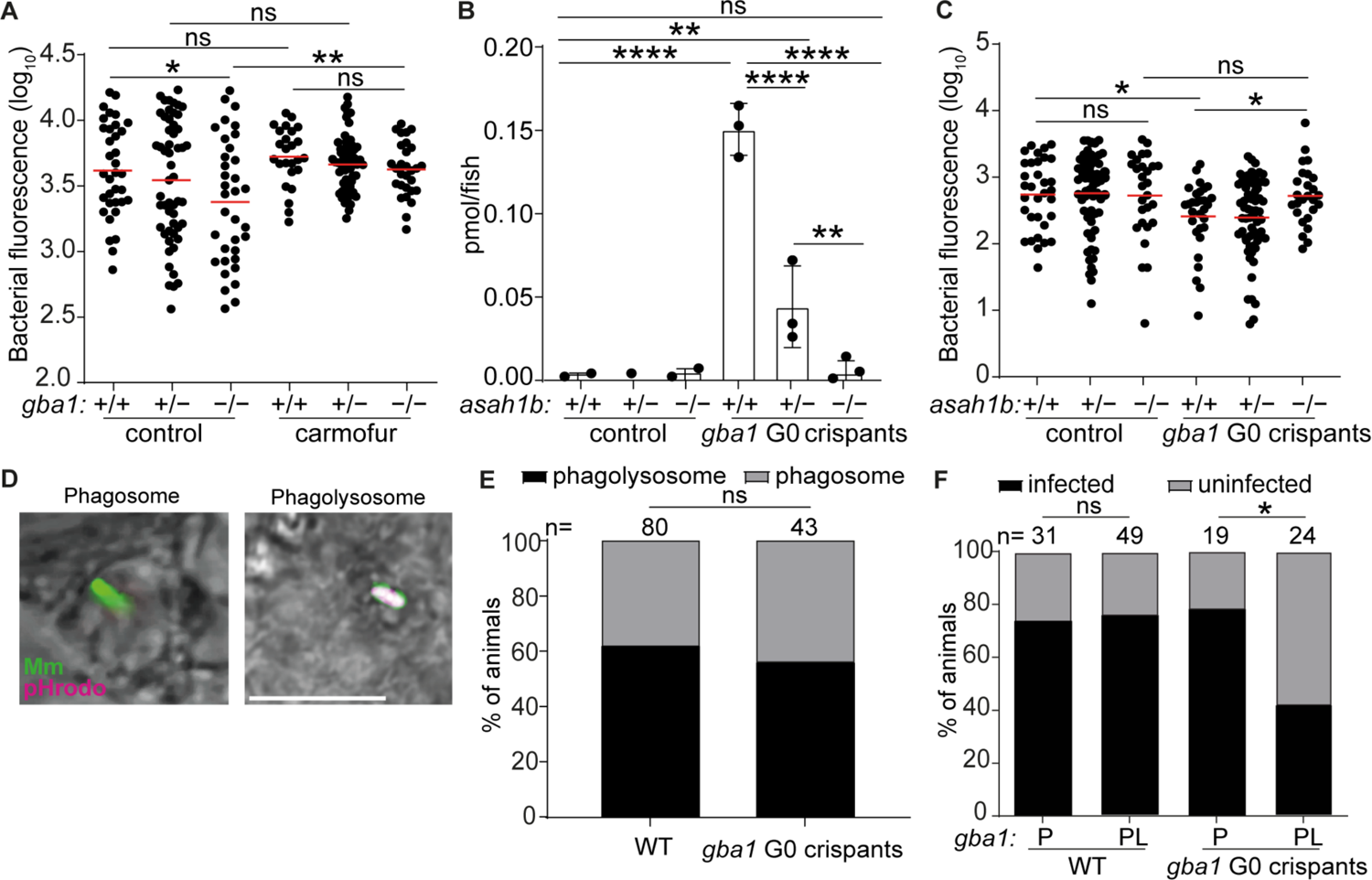
Glucosylsphingosine is bactericidal in vivo. *(A)* Bacterial burden measured by fluorescence in *gba1^sa1621/+^*incross larvae treated with 0.5 μM carmofur or vehicle control 5 dpi after CV infection with 200-300 Mm. Horizontal bars, means; ns, not significant; \**P* < 0.05; \**\*P <* 0.01 (one-way ANOVA with Tukey’s post-test). Representative of 2 independent experiments. *(B)* GlcSph (pmol/fish) in *asah1b* incross larvae that are either wild-type or G0 crispant for *gba1*. Mean+/-S.D; ns, not significant; \**\*P <* 0.01; \*\*\**\*P <* 0.0001 (one-way ANOVA with Tukey’s post-test); Representative of 2 independent experiments. *(C)* Bacterial burden measured by fluorescence in *asah1b* incross larvae that are either wild-type or G0 crispant for *gba1,* 5dpi after CV infection with 200-300 Mm. Horizontal bars, means; ns, not significant; \**P* < 0.05 (one-way ANOVA with Tukey’s post-test). *(D)* Representative confocal images of phagosome-localized Mm (green) and phagolysosome-localized Mm (magenta) at 12 hpi. Scale bar, 20 μm. *(E)* Percentage of animals in which the single infecting Mm bacterium was localized to a phagosome or a phagolysosome; ns, not significant (Fisher’s exact test). Representative of 2-3 independent experiments. *(F)* Percentage of infected wild-type and *gba1* G0 crispants at 5 dpi after HBV infection with a single Mm bacterium. P, phagosome; PL, phagolysosome. ns, not significant; *\*P <* 0.05 (Fisher’s exact test). Representative of 2-3 independent experiments.

To determine if the accumulated glucosylsphingosine in macrophages was also sufficient for resistance in vivo, we took advantage of an important feature of the intra-macrophage lifestyle of mycobacteria, namely that upon infecting macrophages, Mm and Mtb phagosomes frequently but not always fuse to lysosomes (50). Therefore, if glucosyl sphingosine is sufficient for increased macrophage mycobactericidal activity, then only the mycobacteria in the fused phagolysosome compartments should be killed more in GBA-deficient macrophages as compared to wild-type macrophages. If, on the other hand, increased inflammation in GBA-deficient macrophages additionally contributes to the increased killing then mycobacteria in unfused phagosomes should also have a greater likelihood of being killed. By infecting zebrafish larvae with green fluorescent bacteria that have been labeled with pHrodo, a dye that fluoresces red in acidified compartments, we were able to distinguish bacteria in fused phagosome-lysosome compartments (green plus red fluorescence) from those in unfused phagosomes (only green fluorescence) (50) (Fig. 5*D* and *SI Appendix*, Fig. S6). The fate of these individual bacteria – growth versus clearance – can be determined using the infectivity assay at 5 dpi (50) (*SI Appendix*, Fig. S6). We found, as expected, that approximately 60% of the bacteria were found in fused phago-lysosomes in wild-type animals (50); this proportion was not altered in *gba1* mutants (Fig. 5*E*). Also as predicted on account of the acid tolerance of mycobacteria, in wild-type animals phagosome-lysosome fusion did not increase clearance of infection (50) (Fig. 5*F*). By contrast, in *gba1* mutants, bacteria in fused phagosome-lysosomes were significantly more likely to be killed (Fig. 5*F*). However, there was no increased killing of bacteria in the unfused phagosomes of GBA-deficient macrophages (Fig. 5*F*). Thus, the increased bacterial clearance in *gba1* mutants was entirely due to increased microbicidal capacity specifically of their lysosomes (Fig. 5*F*). Therefore, the glucosylsphingosine that accumulates in GBA-deficient macrophage lysosomes is both necessary and sufficient for the increased mycobacterial killing observed in vivo at the early stages of infection, which leads to increased clearing of infection. Indeed, our results show that the subcellular location where bacterial killing occurs is precisely where the formation of glucosylsphingosine is maximal. These findings rule out a role for the increased inflammation associated with Gaucher disease in early resistance and enhanced bacterial clearing.

### The common N370S GBA mutation confers resistance to TB

Our cumulative findings led to the question of whether mutant alleles associated with Gaucher disease protect humans against TB. This question is pertinent to the debate on why the disease incidence persists at high frequency in the Ashkenazi Jewish population (1/800 births versus 1/40,000-60,000 births in other populations) (17). Previous hypotheses of a link between Gaucher disease and protection against TB in Ashkenazi Jews have proposed a model of heterozygote advantage wherein TB resistance in heterozygotes might offset the deleterious effects of the disease in homozygotes (29, 32). However, consistent with glucosylsphingosine accumulation occurring only in homozygotes, we had found that only homozygotes, not heterozygotes, were resistant to Mm (Fig. 2*C*, 2*E* and 2*G*-2*I*). Following up on our finding that glucosylsphingosine also kills Mtb in vitro (Fig. 4*F*), we showed that *gba1* mutant homozygotes were also resistant to Mtb (Fig. 6*A*). Again, heterozygotes remained susceptible (Fig. 6*A*). Thus, our findings support a link between Gaucher disease homozygosity and protection against TB directly through the microbicidal activity of accumulated glucosylsphingosine in macrophage lysosomes, rather than the proposed model of heterozygote advantage presumably through some indirect means.

**Fig. 6.**
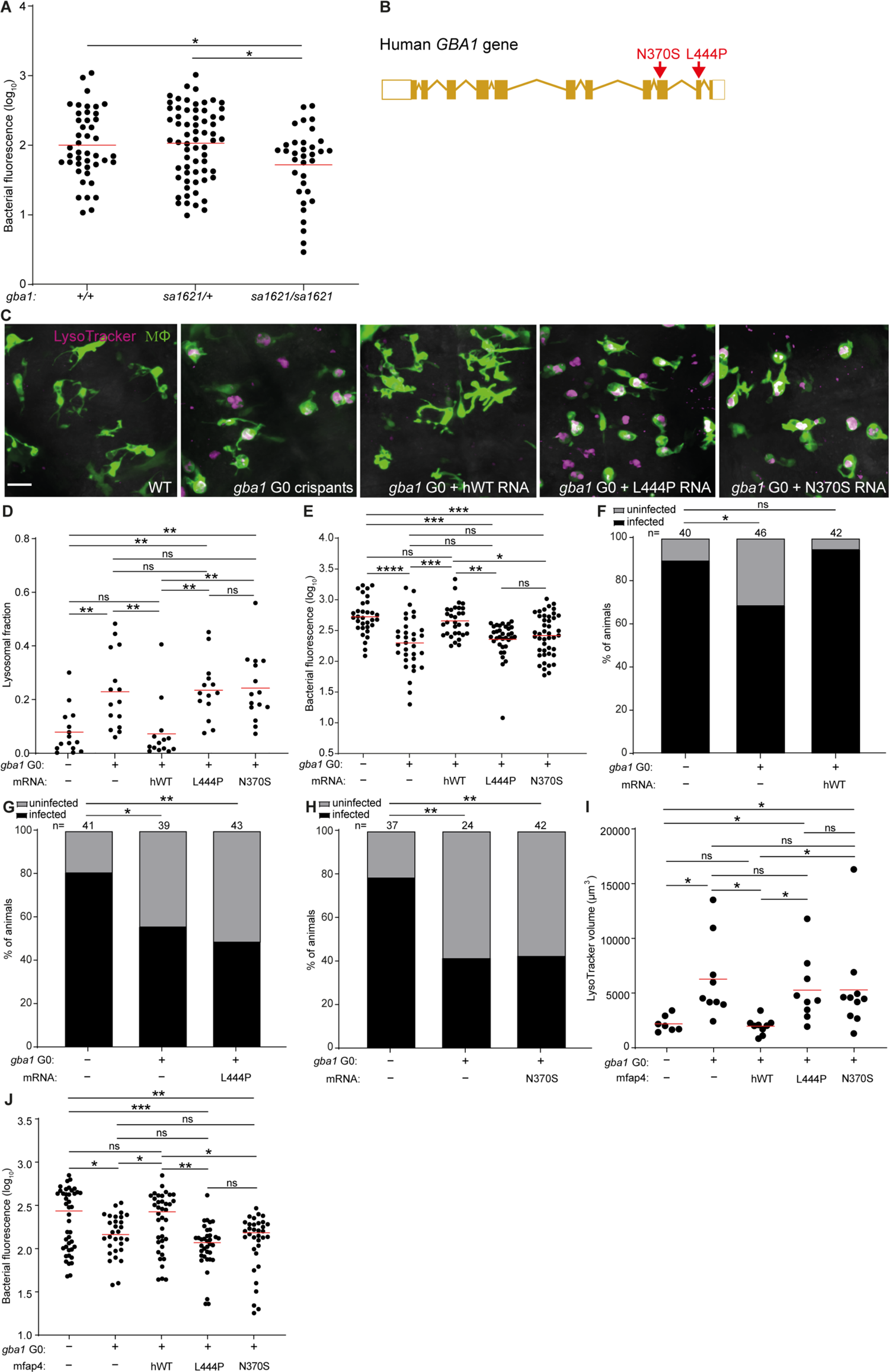
Common human *GBA1* mutants confer resistance to TB. *(A)* Bacterial burden measured by fluorescence in *gba1^sa1621/+^*incross larvae 5dpi after CV infection of 400-500 Mtb. Horizontal bars, means. *\*P <* 0.05 (one-way ANOVA with Tukey’s post-test). *(B)* Schematic diagram showing the location of the two human *GBA1* mutations studied. *(C)* Maximum intensity projection of representative pseudo-colored confocal images of brain macrophages (yellow fluorescent) stained with LysoTracker Red in wild-type larvae, *gba1* G0 crispants, and *gba1* G0 crispants expressing human wild-type, L444P or N370S mutant *GBA1* mRNA. Scale bar, 20 μm. *(D)* Mean LysoTracker red volume per macrophage in 3 dpf animals from *(C)*. Each point represents mean LysoTracker Red volume fraction per macrophage in each fish. Horizontal bars, means; ns, not significant; \**\*P <* 0.01 (one-way ANOVA with Tukey’s post-test). Representative of 2-3 independent experiments. *(E)* Bacterial burden measured by fluorescence in 5 dpi wild-type, *gba1* G0 crispants and *gba1* G0 crispants expressing human wild-type, L444P or N370S mutant *GBA1* mRNA after CV infection with 600-700 Mtb. Horizontal bars, means; ns, not significant; *\*P <* 0.05; \**\*P <* 0.01; \*\**\*P <* 0.001; \*\*\**\*P <* 0.0001 (one-way ANOVA with Tukey’s post-test). Representative of 2 independent experiments. *(F)* Percentage of infected wild-type animals, *gba1* G0 crispants and *gba1* G0 crispants expressing human wild-type *GBA1* mRNA fish at 5 dpi after HBV infection with a single Mtb bacterium. ns, not significant; *\*P <* 0.05 (Fisher’s exact test). Representative of 2 independent experiments. *(G)* Percentage of infected wild-type animals, *gba1* G0 crispants and *gba1* G0 crispants expressing human L444P mutant *GBA1* mRNA at 5 dpi after HBV infection with a single Mtb bacterium. *\*P <* 0.05; \**\*P <* 0.01 (Fisher’s exact test). *(H)* Percentage of infected wild-type animals, *gba1* G0 crispants and *gba1* G0 crispants expressing human N370S mutant *GBA1* mRNA at 5 dpi after HBV infection with a single Mtb bacterium. \**\*P <* 0.01 (Fisher’s exact test). Representative of 2 independent experiments. *(I)* Total LysoTracker red volume in 3 dpf wild-type, *gba1* G0 crispants and *gba1* G0 crispants expressing macrophage-specific human wild-type, L444P or N370S mutant *GBA1* gene. Each point represents total LysoTracker Red volume in each fish. Horizontal bars, means; ns, not significant; *\*P <* 0.05 (one-way ANOVA with Tukey’s post-test). *(J)* Bacterial burden measured by fluorescence in 5 dpi wild-type, *gba1* G0 crispants and *gba1* G0 crispants expressing macrophage-specific human wild-type, L444P or N370S mutant *GBA1* gene after CV infection with 600-700 Mtb. Horizontal bars, means; ns, not significant; *\*P <* 0.05; \**\*P <* 0.01; \*\**\*P <* 0.001 (one-way ANOVA with Tukey’s post-test).

The proposal that homozygosity for a disease-causing allele itself drives positive selection presents a conundrum. This conundrum may be resolved by the widely variable features of disease in patients homozygous for *GBA1* N370S (c.1226 A>G; also referred to as p.Asn409Ser - N409S - to include a signal peptide) (Fig. 6*B*). More than 300 mutations of *GBA1* have been described worldwide (16, 26) but the evidence for selective advantage is centered around the N370S allele, which occurs at high frequency in the Ashkenazi Jewish population, with between 1 in 18 and 1 in 11 persons a carrier of the mutation (25–28). Persons homozygous for GBA N370S typically manifest the mildest (non-neuronopathic, Type 1) form of Gaucher disease with age of onset 10-30 years later than for other Gaucher genotypes (25, 51). Indeed, half to two-thirds of individuals homozygous for GBA N370S remain asymptomatic, and fertility is normal even among homozygotes with Gaucher disease-associated pathologies (16, 25, 52). These characteristics of N370S Gaucher disease may be consistent with a model where protection of homozygotes against TB offsets the deleterious effect of the mutation in the Ashkenazi population as a whole.

For the homozygote protection model for GBA N370S to be plausible, two essential conditions must be met: 1) glucosylsphingosine must accumulate at mycobactericidal concentrations in GBA N370S homozygote macrophages, similar to the case in GBA-deficient zebrafish macrophages; and 2) zebrafish expressing the human GBA N370S mutation must be resistant to Mtb, similar to zebrafish GBA null mutants. To see if the first condition was met, we calculated glucosylsphingosine concentrations in macrophages from published reports of amounts determined in macrophages prepared from humans homozygous for the GBA N370S mutation as well as from healthy human macrophages treated with a glucocerebrosidase inhibitor; each are reported to be ≈0.6 nmol/mg protein (18, 39) (*SI Appendix*, Supplementary Box S1). We calculated these intramacrophage concentration to be ≈200 μM (*SI Appendix*, Supplementary Box S1). This is in excellent agreement with glucosylsphingosine concentrations of 19 μM (95% CI 9-29) in whole spleen tissue obtained from N370S compound heterozygotes with Type 1 non-neuronopathic Gaucher disease (*SI Appendix*, Supplementary Box S1), in which the lipid-laden macrophages or Gaucher cells accumulate but still contribute only a minor fraction of tissue mass (18, 39). Thus, the intramacrophage concentration of glucosylsphingosine exceeds its MIC for Mtb (60 μM), and as a base the molecule will be further concentrated in the acidic lysosome compartment of macrophages as positively charged micelles with potentiated bactericidal activity (53, 54).

To test the second condition that must be met, namely whether the human GBA N370S mutation results in resistance to Mtb, we turned to genetic complementation of the zebrafish *gba1* G0 crispants with human *GBA* RNAs, which have been used previously to rescue bone abnormalities in *gba1* mutant zebrafish (20). Wild-type human *GBA* RNA should reverse their macrophage lysosomal storage phenotype as well as eliminate Mtb resistance (as shown with zebrafish *gba1* RNA in Mm (Fig. 2*J*). In contrast, neither of the two phenotypes should be altered by RNA bearing the L444P mutation (c.1448 T>C and also referred to as p.Leu483Pro, L483P), a severe Gaucher disease allele with minimal residual glucocerebrosidase activity in vitro (26, 55). As predicted, injection of wild-type *GBA* RNA, but not L444P *GBA* RNA, rescued lysosomal storage and abolished Mtb resistance, demonstrating the validity of this complementation assay (Fig. 6*C*-6*E*). We next tested whether the zebrafish *gba1* mutant phenotypes would be reversed by the human N370S GBA allele. Despite its greater residual glucocerebrosidase activity (10-15% of wild-type) (56), N370S *GBA* RNA also failed to rescue lysosomal storage and abolish Mtb resistance (Fig. 6*C*-6*E*). To test if the N370S GBA mutation renders Mtb-infected macrophages microbicidal, we performed the infectivity assay. Wild-type *GBA* RNA restored the reduced infectivity of the *gba1* G0 crispants to wild-type levels whereas neither mutant RNA did (Fig. 6*F*-6*H*).

The experiments above used transient, ubiquitously expressed human RNAs to evaluate human GBA alleles for protection against TB. In an independent approach, we created stable transgenic zebrafish lines expressing either wild-type or each of the two mutant GBA alleles selectively in macrophages (*SI Appendix*, Fig. S7). GBA1 crispants were created either in wild-type or in each of the three transgenic backgrounds, resulting respectively in GBA1 null mutants, or mutants expressing human wild-type, L444P or N370S GBA1, but only in their macrophages. As was the case with the transiently expressed human RNAs, macrophage-specific expression of human wild-type GBA1 reversed both the macrophage lysosomal phenotype and the resistance to TB of the GBA null mutants whereas neither the L444P nor the N370S GBA alleles did (Fig. 6*I* and 6*J*). We could not test the infectivity phenotypes in this set of experiments because of the paucity of available transgenic mutants - ∼ 5-10 fold more animals are required to obtain sufficient animals who received single bacteria; however, since this is a macrophage-intrinsic phenotype, there is no reason for the results to be different from those obtained for the animals complemented with the ubiquitously-expressed RNAs. Thus, in the zebrafish, the human N370S GBA allele has the same effect as more severe Gaucher disease mutations, resulting in lysosomal storage and conferring increased resistance to TB through increased macrophage microbicidal activity against Mtb.

## Discussion

Our findings provide support for the longstanding hypothesis that Gaucher disease, including the common Ashkenazi Jewish N370S mutation, confers resistance to TB by enhancing macrophage mycobactericidal activity. We find that in Gaucher disease, macrophage mycobactericidal activity is enhanced from the earliest step of infection when macrophages first encounter mycobacteria. In the context of the Mtb life-cycle, this would occur when alveolar macrophages in the lung phagocytose the bacteria (1). Enhanced macrophage mycobactericidal capacity would also increase the chances of clearing infection in the granuloma. For macrophages to be recruited to and become infected with mycobacteria they must be motile, which is unlikely the case for the classic Gaucher cell with substantial lysosomal storage (7, 18). However, this work shows that even macrophages without obvious lysosomal storage have increased microbicidal capacity, leading us to infer that the macrophages in Gaucher disease that mediate resistance have accumulated smaller amounts of glucosylsphingosine, enough to restrict mycobacterial growth, though not enough to induce the Gaucher cell phenotype. In support of this, while alveolar macrophages can manifest lysosomal storage in Gaucher disease, frank Gaucher cells within airspaces are a rare occurrence in patients with type 1 disease – the non-neuronopathic phenotype of N370S homozygotes (57–59). Our estimate that even 15 μM concentrations of glucosylsphingosine - far lower than the ∼200 μM concentrations found in GBA-deficient macrophages – inhibits mycobacterial growth lends further support to this argument. Macrophages with this much lower glucosylsphingosine concentration may well lack the classical Gaucher cell phenotype but possess enhanced ability to restrict mycobacteria, particularly given the concentration of the compound in the form of positively charged micelles in mycobacterial phagolysosomes, which we have demonstrated to be mycobactericidal in GBA-deficient zebrafish.

This work highlights the advantages of the zebrafish with its amenability to genetic and pharmacological manipulations and the unique live subcellular imaging possible in this transparent model organism. Particularly germane to this study, GBA-deficient macrophages will have different lipid loads at different times, depending on when they have ingested dying cells (7). The use of the zebrafish has allowed us to study GBA-deficient macrophages, including those carrying the human N370S mutation in a true in vivo context, while in the process of carrying out their dual homeostatic scavenger and anti-microbial functions.

Our findings have potentially important genetic implications. They offer biological evidence related to the question of whether GBA N370S has persisted at high frequency because it conferred a selective advantage or solely because of multiple, severe population bottlenecks (29–32). There has been debate over the decades about how a disease allele could persist at a relatively high frequency in the Ashkenazi Jewish population when it is obviously detrimental (29, 60). The observation that other genetic lysosomal disorders (e.g., Tay-Sachs, Niemann-Pick) are also found at elevated frequency in this population suggested the possibility of positive selection of these detrimental alleles, possibly from a heterozygote advantage (29, 30, 33, 60, 61).

TB has been cited as the potential selective force, amid speculation that densely populated urban living conditions in ghettoes might have placed Jews under stronger selection than other Europeans (29, 30, 60–62). TB was a significant killer of young people in Europe through the Middle Ages into the 19^th^ century (63, 64). Indeed, an estimate of mortality based on historical records from 1891-1900 attributed half of all deaths in the reproductive years (ages 25-40) to TB, calculated to lead to a loss of 7-15% of reproductive fitness per generation compared to a hypothetical cohort of completely TB-resistant individuals (63). It is difficult to determine the exact age of the N370S allele owing to confounding variables such as admixture (65, 66), uncertain mutation and recombination rates, and conflicting data on historical population sizes (32). Nevertheless, all estimates based on haplotype analyses agree that it is at least 800 years old, with some finding it to be as much as 1400 years old (32, 67, 68). Therefore, N370S would have been present in the Ashkenazim through the centuries when their exposure to TB would have been high. If individuals homozygous for GBA N370S were more resistant, they would have had a substantial survival and reproductive advantage during periods of high TB transmission.

In keeping with biallelic inheritance where only homozygotes and not heterozygotes accumulate the culpable substrate, only homozygotes and not heterozygotes are resistant to TB. We argue therefore for a model of homozygote protection against TB as a selective force, rather than the generally accepted “balanced polymorphism” concept of heterozygote advantage offsetting the deleterious effects of homozygosity (29–32, 69). This model is entirely plausible because homozygous N370S Gaucher disease is relatively mild (2/3 of N370S homozygotes are asymptomatic) (29, 51) and therefore it is compatible with reproductive success. Homozygosity for the N370S allele is associated with a much later age of onset, and fertility is normal even in those individuals who are symptomatic of Gaucher Disease-associated pathologies (16). Given glucosylsphingosine’s broad antibacterial activity, it is intriguing to speculate that GBA N370S homozygotes might have been protected against common but consequential bacterial infections other than TB, contributing additionally to the selective advantage of this allele.

## Materials and Methods

### Zebrafish husbandry

Zebrafish husbandry and experimental procedures were conducted in compliance with guidelines from the UK Home Office. All zebrafish were maintained in buffered reverse osmotic water systems and were exposed to a 14-hour light −10-hour dark cycle to maintain proper circadian conditions. They were fed twice daily a combination of dry food and brine shrimp. The *gba1* mutant fish were culled at 2.5 months old as soon as clinical signs of disease were observed.

### Fish Lines

The mutant lines *gba1^sa1621^* and *asah1b^sa19461^* were created by ENU (N-ethyl-N-nitrosourea) mutagenesis as part of The Zebrafish Mutation Project (ZMP) (70) and provided by the Wellcome Sanger Institute. The allele *gba1^sa1621^* has been previously used to study larval bone ossification (71). The *gba1^sa1621^*and *asah1b^sa19461^* lines were generated in the Tupfel long-fin (TL) strain and outcrossed to the AB wild-type strain (Zebrafish International Resource Center) to be maintained in a mixed AB/TL background. The *gba1^cu41^*and *gba1^cu42^* mutant lines and the transgenic lines *Tg(mpeg1:YFP)^w200^*(with YFP-expressing macrophages) (10) and *Tg(mpeg1:Brainbow)^w201^* (with tdTomato-expressing macrophages) (72) were created and maintained in the AB background. The transgenic zebrafish lines *Tg(BH:eGFP-mfap4:hWT)^cu57^*, *Tg(BH:eGFP-mfap4:N370S)^cu58^*, and *Tg(BH:eGFP-mfap4:L444P)^cu59^*, were generated and maintained as described before (11); briefly, the human wild-type *GBA1* cDNA (hWT) (Sino Biological, NM_000157) was amplified by Phusion High-Fidelity (NEB) PCR. The 3’ adenine (A) overhangs were then added by Taq (NEB) PCR and cloned into the TOPO-TA plasmid (Thermo Fisher Scientific). The human wild-type *GBA1*-containing TOPO-TA plasmid was then used as a template to generate the N370S and L444P *GBA1* variants, using the Q5^®^ Site-Directed Mutagenesis Kit (NEB). Then, each version was inserted after the macrophage-specific mfap4 promoter into the plasmid pTol2-PhiC31LS-BH:GFP-mfap4:New MCS and these plasmids were used to generate the transgenic lines (11). G0 *gba1*-deficient zebrafish embryos (crispants) were generated using CRISPR-Cas9 technology by simultaneously targeting different sites of the *gba1* gene (73). Guide RNAs were prepared following the manufacturer specifications, hybridizing the common RNA component (Alt-R tracrRNA) with each of the specific Alt-R crRNA: crRNA1 (5’-CATACTGCGACTCTCTCGGC-3’) crRNA3 (5’-GAGTGCTGGACCTAGATCCA-3’) crRNA4 (5’-GGGTGACGTCAGGCGGCACA-3’) 3-5 nl of a solution containing Alt-R crRNA and Alt-R tracrRNA (30 μM each) complexed with Cas9 protein (0.25 μg/μl) (Integrated DNA Technologies), and 2% phenol red sodium salt (Sigma) was injected into 1-2 cell stage embryos (73). Similar volumes of a solution containing Cas9 protein and phenol red was used to generate the control animals. The genotype of individual larvae and mutagenesis efficacy were assessed by high-resolution melt (HRM) analysis (74) using the following primers: *gba1* G0 Forward-1 (5’-GGTCATGGCTCGGTTGTGT-3’) *gba1* G0 Reverse-1 (5’-CTCCATCAGTCTGCTGCCAG-3’) *gba1* G0 Forward-3 (5’-AGGCTTTGGGTTTTACTCCG-3’) *gba1* G0 Reverse-3 (5’-TCATTAGTTGGGTCTTGGAAAAG-3’) *gba1* G0 Forward-4 (5’-CACGATATGTCCACGGCATT-3’) *gba1* G0 Reverse-4 (5’-GGAAGTAATCAGGGTACAGATGGT-3’) The mutant lines *gba1^cu42^*and *gba1^cu41^* were generated by targeting the *gba1* gene with Alt-R crRNA3 (the same crRNA3 as used for the G0 crispants) and Alt-R crRNA5 (5’-GGATCGTGGTGTGCGTCT-3’). Genotypes were determined by high-resolution melt analysis (HRM) (74) using the primers *gba1* G0 Forward-3 and *gba1* G0 Reverse-3 for *gba1^cu42^* and *gba1^cu41^* Forward-5 (5’-CCATCTGTACCCTGATTACTTCC-3’), *gba1^cu41^*Reverse-5 (5’-TTAAGACCATTATCCAGTACCTGG-3’) for *gba1^cu41^*. Founders were identified in the F1 generation and the lines were outcrossed to the AB wild-type strain.

### Bacterial strains

*Mycobacterium marinum* (Mm) M strain (ATCC #BAA-535) expressing tdTomato (red fluorescence), mWasabi (green fluorescence) or EBFP2 (blue fluorescence) under the *msp12* promoter (75) were grown under hygromycin (Mediatech) selection in Middlebrook 7H9 medium (Difco), supplemented with oleic acid-albumin-dextrose-saline (OADS), and Tween-80 (Sigma) (75) at 33°C. *Mycobacterium smegmatis* (Msm)(mc^2^ 155) was grown in Middlebrook 7H9 medium supplemented with albumin-dextrose-catalase (ADC) at 37°C (76). *M. tuberculosis* (Mtb) H37Rv *ΔleuD ΔpanCD* double auxotroph (mc^2^ 6206) expressing tdTomato under the *msp12* promoter was grown under hygromycin B selection in Middlebrook 7H9 broth supplemented with oleic acid-albumin-dextrose-catalase (OADC), Tween-80 (Sigma), 0.05 mg/ml L-leucine and 0.024 mg/ml calcium pantothenate (Sigma) at 37°C (11, 77). Single cell suspensions were prepared of all mycobacterial strains to infect zebrafish and for MIC/MBC tests.(75). *Staphylococcus aureus* (ATCC 29213) and *Escherichia coli* (ATCC 25922) were passaged twice on tryptic soy agar with 5% sheep’s blood (Thermo Scientific) at 37° C in 5% CO_2_ for 18-24 hours. 3 to 5 isolated morphologically normal colonies selected from the agar plates were suspended in cation adjusted Mueller Hinton broth (CAMHB, Sigma-Aldrich). The bacterial suspensions were diluted with normal saline to achieve a turbidity equivalent to a 0.5 McFarland standard (latex, Thermo Scientific) using either a Wickerham card (Thermo Scientific) or by nephelometry (DensiCHEK Plus, bioMérieux) before drug exposure.

### Assessment of clinical signs of disease in adult *gba1* deficient zebrafish

As soon as clinical signs of disease were observed, photographs and videos of adult *gba1* zebrafish were taken using the camera incorporated within a cell phone (Huawei P10). Immediately thereafter, the animals were euthanized by tricaine overdose followed by brain destruction according to Schedule 1 of the Animals (Scientific Procedures) Act 1986. Genotypes were confirmed post-mortem.

### Assessment of drug minimum inhibitory concentrations (MIC) and minimum bactericidal concentrations (MBC)

Glucosylsphingosine (Sigma-Aldrich) and D-erythro-Sphingosine (C18) (Santa Cruz Biotechnology) were dissolved in absolute ethanol (Honeywell) at 6 mM. Glucosylceramide (GlcCer, Sigma-Aldrich) was dissolved in DMSO-ethanol (1:1, v/v) at 20 mM. Rifampicin (Sigma) was dissolved in methanol (Honeywell) at 25 mg/ml. Aliquoted drugs were stored at −20°C and used within a month.

All MIC/MBC tests were performed according to CLSI instructions (M07-11^th^ edition for broth MICs and M26A-1999 for bactericidal determination) (78, 79) with modifications as specified below for mycobacterial testing. On the day of the MIC/MBC assessment, all drugs were diluted in broth media twofold to achieve the desired test concentrations. Liquid broth containing 4% absolute ethanol or 4% DMSO-ethanol (1:1, v/v) showed no inhibition of bacterial growth. 100 μl of liquid broth containing varying concentrations of each drug were added in duplicate into sterile round bottom polystyrene 96-well plates (Corning). The inocula for *S. aureus* (ATCC 29213) and *E. coli* (ATCC 25922) were prepared by making a 1:100 dilution in CAMHB of the 0.5 McFarland standard equivalent turbidity of the bacterial culture. Mycobacterial inocula were prepared by diluting single cell suspensions to OD_600_ 0.03. 100 μl of bacterial culture (without Tween-80) was added to the previously prepared plate containing 2X final drug concentrations. The final bacterial inoculum of approximately 5 x10^5^ cfu/ml in the MIC/MBC test was confirmed by counting CFU on agar plates.

Drug-treated *S. aureus* and *E. coli* microbroth plates were incubated at 37°C (5% CO_2_) for 16-20 hours without shaking. Drug-treated Mm microbroth plates were incubated at 33°C for 5-7 days without shaking. Drug-treated Msm was incubated at 37°C for 2-3 days without shaking. Drug-treated Mtb was incubated at 37°C for 14-21 days in 5% CO_2_ without shaking. The MIC was defined as the lowest concentration of drug which completely inhibited bacterial growth as detected by eye. According to CLSI instructions, the MIC of the quality control drug rifampicin should be: 0.004 to 0.016 mg/L for *S. aureus*, and 4 to 16 mg/L for *E. coli*. The MBC was defined as a ≥3log_10_ decrease from the initial concentration by plating drug treated bacterial culture with ten-fold serial dilutions after the MIC was determined. *S. aureus* and *E. coli* were plated on CAMHB media at 37°C overnight. Mm was plated on 7H10+OADS media at 33°C for 7-9 days. Msm was plated on 7H10+ADC media at 37°C for 2-3 days. Mtb was plated on 7H10+OADC+leu+pan media at 37°C and grown in a humidified incubator with 5% CO_2_ for a month. Drug carryover inhibiting bacterial growth on the culture plates was excluded because the ten-fold dilutions resulted in drug concentrations below their respective MICs for the test organisms.

### Zebrafish infections

Zebrafish embryos were housed in fish water (reverse osmosis water containing 0.18 g/l Instant Ocean supplemented with 0.25 mg/ml methylene blue) up to 1 day post-fertilization (dpf) at 28.5°C. From 1 dpf, embryos were maintained in 0.5X E2 medium (80) with 0.003% PTU (1-phenyl-2-thiourea, Sigma).

To assess bacterial burdens, zebrafish were infected with Mm at 2 dpf in the hindbrain ventricle (HBV) or the caudal vein (CV) (75). Inocula of 200–300 and 100– 150 Mm were used respectively, for CV and HBV. Mtb bacterial burdens were assessed after CV infection using inocula of 400–800 bacteria. For infections using a heterozygotes incross to generate all genotypes, larvae were genotyped using KASP (*gba1^sa1621^* and *asah1b^sa19461^*) or HRM (*gba1^cu41^* and *gba1^cu42^*) assays at the end of experiment.

For infectivity assays, highly diluted mWasabi-expressing Mm (0-3 cfu/nl) or tdTomato-expressing Mtb (0-3 cfu/nl) were injected into the HBV at 2 dpf. Staining with pHrodo Red to assess phagosomal versus phagolysosomal localization of the bacteria was performed before infection into the HBV as described (50). The phagosome- and phagolysosome-located Mm were sorted 12-24 hours post infection by confocal microscopy.

## Microscopy

### Assessment of macrophage morphology and speed

To assess macrophage vacuolation, 1 mM LysoTracker Red DND-99 (Life technologies) in DMSO solution was diluted 1:25 in PBS and injected into the CV of 3 dpf zebrafish larvae. Larvae were then incubated at 28.5°C in the dark for 1 hour before confocal imaging. To assess engulfment accumulation of cell debris in macrophages, acridine orange (ImmunoChemistry, Bloomington, MN) was dissolved in 0.5X E2 fish water at 2 μg/mL and larvae were soaked in this solution at 28.5°C in the dark for 30 minutes before imaging. Macrophage morphology was assessed using a Nikon A1R confocal microscope with a 20X Plan Apo 0.75 NA objective with a galvano scanner used to generate 40 μm *z*-stacks consisting of 1.3–2 μm optical sections. Images were acquired and processed with NIS Elements (Nikon) using the Denoise.ai function and maximum intensity projection. Macrophage, LysoTracker and acridine orange volumes were measured using the surface rendering feature of Imaris 9.1 (Bitplane Scientific Software).

For macrophage speeds, using the same settings as above, timelapse images were taken using a resonant scanner at 3-minute intervals for 2 hours maintaining the larvae at 28.5°C using a heating chamber (Okolab). Macrophage tracks were generated by surface rendering and object classification features of Imaris 9.7 (Bitplane Scientific Software), and the mean speed of each track calculated. When the fish used in experiments were generated from *gba1* heterozygote incrosses, their genotype was determined by KASP or HRM assay at the end of experiment, ensuring blinded quantification and analysis.

### Assessment of infection

Bacterial burdens were assessed 3 dpi for HBV infections and 5 dpi for CV infections. To assess bacterial burdens in HBV infections, zebrafish larvae were anesthetized in 0.5X E2 media containing 0.025% Tricaine and embedded in 2.0% low melting point agarose on 6-well optical bottom plates (MatTek Corporation) before imaging. For CV infections, animals were anesthetized in 0.025% tricaine (Sigma-Aldrich) before imaging. Bacterial burdens were assessed by fluorescence pixel counts (FPC) using wide-field microscopy was performed using a Nikon Eclipse Ti-E equipped with a C-HGFIE 130W LED light source and 4X objective. Fluorescence images were captured with a CoolSNAP HQ2 Monochrome camera (Photometrics) using NIS-Elements (version 3.22). Fluorescence filter cubes sets included Chroma FITC (41002), Cy3/TRITC (41004), for detection of green and red fluorescence, respectively.

For HBV infection, Mm volume was assessed using a Nikon A1R confocal microscope with a 20X Plan Apo 0.75 NA objective with a galvano scanner used to generate 40 μm *z*-stacks consisting of 1.3–2 μm optical sections. Images were acquired and processed with NIS Elements (Nikon) using the Denoise.ai function and maximum intensity projection. After HBV infection, bacterial volumes were measured using the surface rendering feature of Imaris 9.1 (Bitplane Scientific Software).

For the infectivity assays, as soon as all the animals had been infected, larvae that had been infected with a single bacterium were identified as follows: the larvae were gently embedded in 3% methylcellulose in 6-well optical bottom plates (MatTek Corporation) in a supine position and the number of infecting bacteria counted using a Nikon Eclipse Ti-E inverted microscope with a 10 or 20X objective and 10X ocular. At 5dpi, the larvae were assessed for the presence or absence of bacteria using a Nikon Eclipse Ti-E inverted microscope with a 10X or 20X objective and 10X ocular.

### Drug administration to zebrafish larvae

Carmofur (Abcam) was dissolved in DMSO (Sigma) and stored in small aliquots at 50 mM at −20°C. 1 dpf embryos were treated fish water with 0.5 μM carmofur (Abcam) with the corresponding volume of 1% DMSO used for the control group. The fish water was renewed daily with freshly prepared drug until the end of the experiment.

To administer VPRIV® (velaglucerase alfa), 2 dpf larvae infected in the HBV were randomly distributed into control and velaglucerase-treated groups at four hours post-infection and 1-1.5 nl PBS containing velaglucerase alfa (100 units/ml) or PBS alone injected into the HBV.

### Zebrafish and human mRNA expression in zebrafish

Total RNA was extracted from 14 dpf wild-type zebrafish and reverse transcribed into cDNA with PrimeScript™ 1st strand cDNA Synthesis Kit (Takarabio). The zebrafish wild-type cDNA and human wild-type *GBA1* (Sino Biological, NM_000157) were amplified by Phusion polymerase (NEB) PCR with a forward primer containing the sequence for the T7 promoter (5’-TAATACGACTCACTATAGG-3’) followed by a Kozak sequence (5’-GCCGCCACC-3’). The 3’ adenine (A) overhangs were then added by Taq PCR (NEB) and cloned into the TOPO-TA plasmid (Thermo Fisher Scientific) and cloned into the TOPO-TA plasmid (Thermo Fisher Scientific).

The human wild-type *GBA1* TOPO-TA plasmid was used as a template to generate the N370S and L444P *GBA1* variants, using the Q5^®^ Site-Directed Mutagenesis Kit (NEB). Primers designed with 5’ ends annealing back-to-back (NEB online design software NEBaseChanger™) were used to produce all *GBA1* variants. The primer sequences are: N370S-Q5-F (5’-ATCATCACGAgCCTCCTGTAC-3’) N370S-Q5-R (5’-GCTGTGGCTGTACTGCAT-3’) L444P-Q5-F (5’-AAGAACGACCcGGACGCAGTG-3’) L444P-Q5-R (5’-CTGACTGGCAACCAGCCC-3’) The mMESSAGE mMACHINE™ T7 Transcription Kit (Thermo Fisher Scientific) followed by Poly(A) Tailing Kit (Thermo Fisher Scientific) was used for *in vitro* mRNA production. 3-5 nL of a solution containing each RNA (200 ng/ul and 300 ng/ul for zebrafish and human, respectively), 0.5X Tango Buffer (Thermo Scientific), and 2% phenol red sodium salt (Sigma) was injected into the yolk of 1-2 cell-stage embryos.

### Negative stain-transmission electron microscopy (TEM)

For TEM experiments, bacteria were treated with drugs as for the MIC/MBC experiments. CFU counts were determined by plating before and after drug treatments. Bacterial cultures were spun down and gently washed twice with a 10 mM Tris–HCl solution containing 5 mM glucose (MRC LMB Media and Glass Wash) and a final concentration of 2 x 10^7^-2 x10^8^ cfu/ml bacteria (OD_600_=0.1-1) was used to prepare the electron microscopy grids.

For negative staining, carbon-coated 300-mesh electron microscopy grids (Agar Scientific) were glow discharged just prior to use (PELCO easiGlow; 45 seconds at 15-25 mA). Grids were incubated for 1 minute with 3 μL bacterial culture, prepared as above, then side-blotted to remove excess. A 3 μL drop of 0.75% uranyl formate was applied to the grid and immediately side-blotted, before the addition of a second 3 μL drop of 0.75% uranyl formate. This was incubated for 30 seconds before side blotting as before. Grids were allowed to dry completely before TEM imaging. TEM imaging was carried out at 120 keV on a ThermoFisher (Formerly FEI) Tecnai G2 Spirit equipped with Gatan Orius SC200W CCD camera. For quantification, grid squares with good staining were randomly selected and every cell in those squares was imaged.

### (Glyco)sphingolipid analysis

Lipids were extracted and analyzed from 5 dpf individual zebrafish larvae as described previously (21, 81).

### Assessment of Gba1 activity

β-glucocerebrosidase (GCase) activity was measured as described earlier (82), using homogenates prepared from pools of 5 dpf zebrafish (10 zebrafish larvae per sample) in potassium phosphate lysis buffer containing: 25 mM K_2_HPO_4_-KH_2_PO_4_ pH 6.5, 0.1% (v/v) Triton-X100 and EDTA-free protease inhibitor (cOmplete™, EDTA-free Protease inhibitor Cocktail, Roche, Sigma-Aldrich). All activities were measured using three independent homogenates in technical duplicate.

### Statistical Analysis

The following statistical analyses were performed using Prism 9 (GraphPad): Student’s unpaired t-test, one-way ANOVA with Tukey’s post-tests and Fisher’s exact test. \**P <* 0.05; \*\**P <* 0.01; \*\*\**P <* 0.001; *P* < 0.0001; not significant, *P* ≥ 0.05.

## Acknowledgments

We thank M.-C. King for encouragement, advice, and critical review of the manuscript, A. J. Pagán for technical guidance and manuscript review, N. Yamaguchi for assistance with infectivity assays and manuscript review, J. Löwe and M. Behr for manuscript review, K.K. Takaki for assistance with infectivity assays and help and guidance on figure preparation, J. K. Shanahan for discussion and assistance with infectivity assays, A. Fountain for discussion and N. Goodwin and other staff of the University of Cambridge aquatics facility for zebrafish husbandry and assistance with genotyping. This work was supported by a Wellcome Trust Principal Research Fellowship (223103/Z/21/Z) and NIH MERIT award (R37 AI054503) to L.R., a Gates Cambridge scholarship to J. F., an NWO grant (BBOL-2007247202) to J. M. F. G. A., National Institute for Health Research UK Cambridge Biomedical Research Centre grant (IS-BRC-1215-20014) to T.M.C. and the MCIN and “ESF Investing in your Future” RYC2019-027799-608 I/AEI/10.13039/501100011033 fellowships to F.J.R since September 2021.

## SI Appendix

### Supporting Information

##### Supplementary Box S1

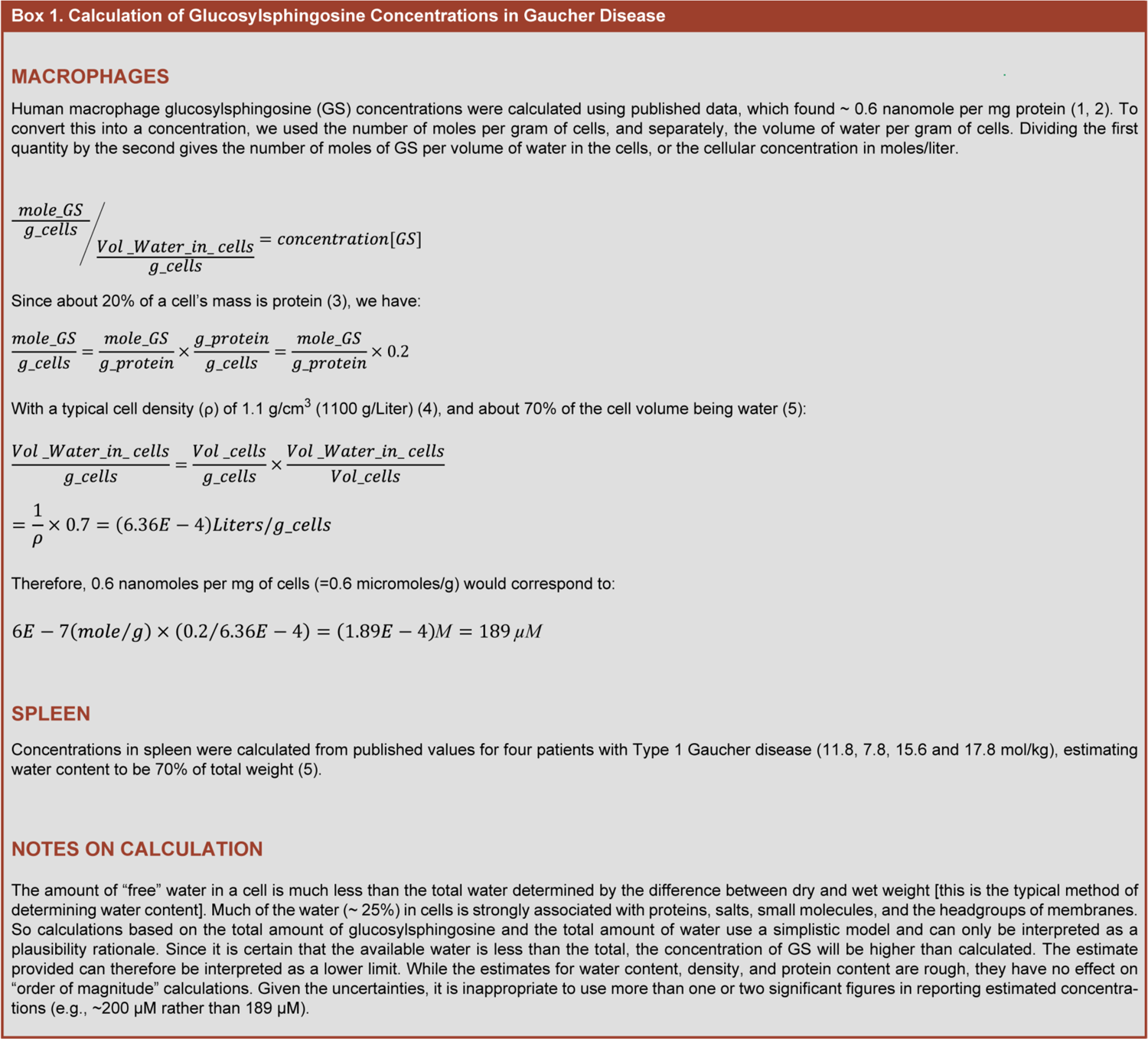

**Fig. S1.**
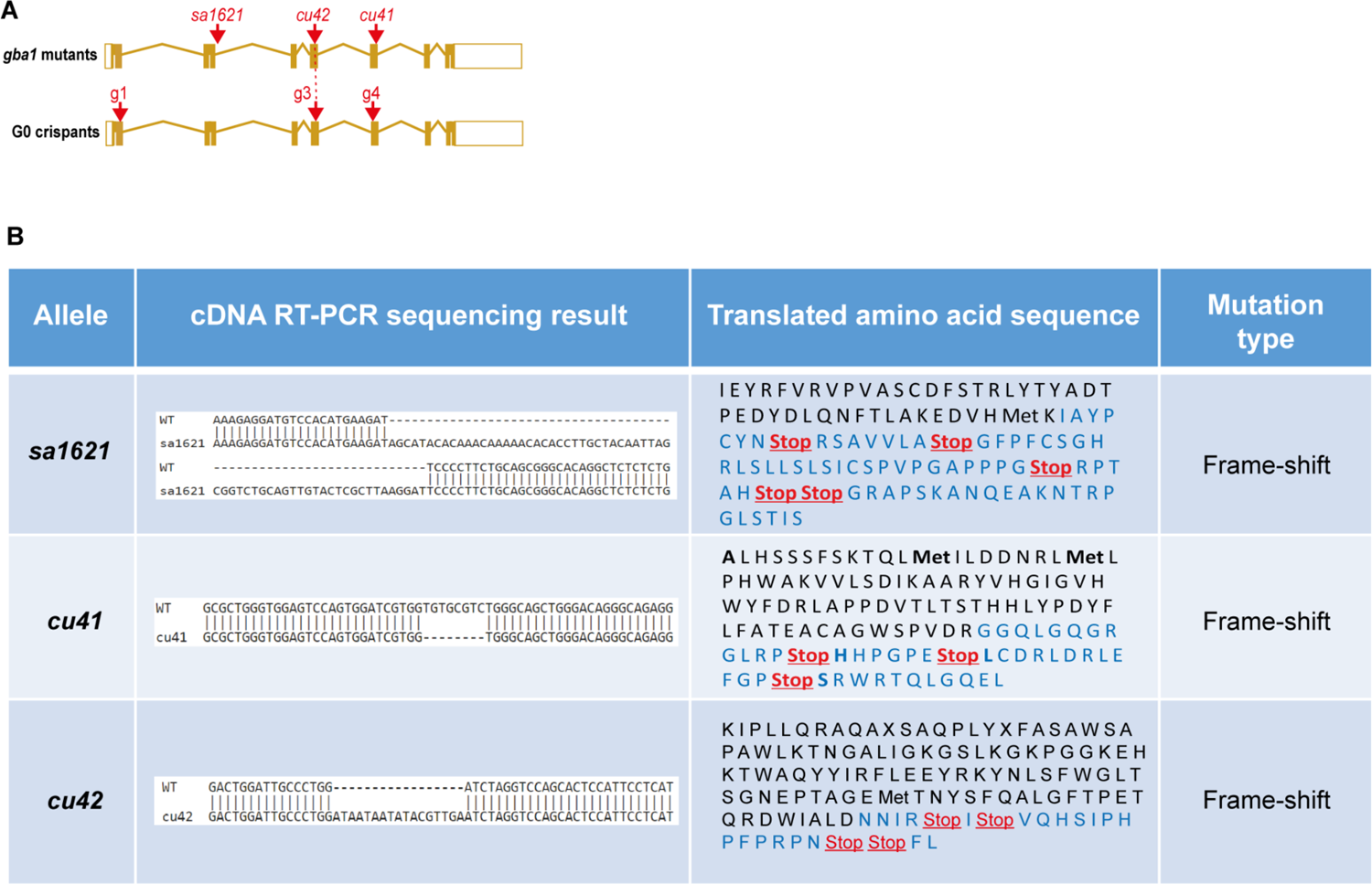
*gba1* mutations characterization. *(A)* Diagram of the zebrafish *gba1* gene showing the locations of the three zebrafish mutations used in this study. *(B)* cDNA sequences of the three mutations and predicted amino acid sequence.

**Fig. S2.**
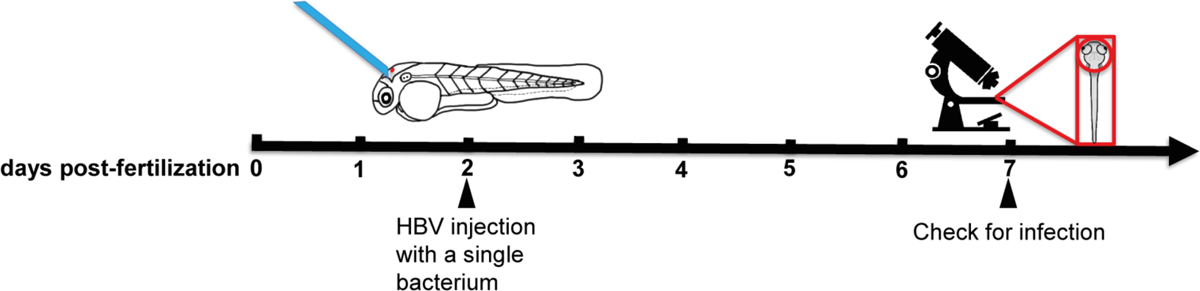
Schematic diagram of the infectivity experiment.

**Fig. S3.**
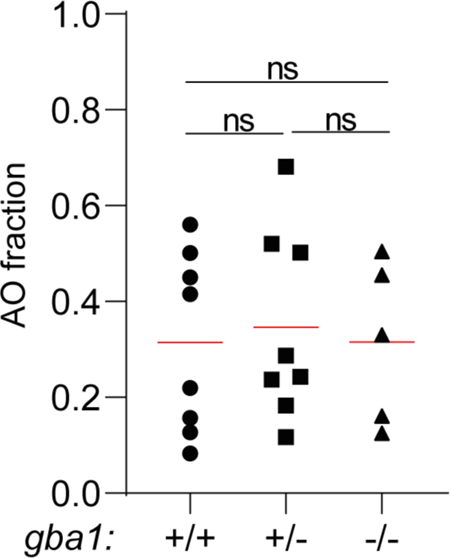
Quantification of AO volume per macrophage in the caudal hematopoietic tissue (CHT) of 5 dpf zebrafish. Each point represents the average AO volume fraction per macrophage in each fish. Horizontal red bars, means. ns, not significant (one-way ANOVA with Tukey’s post-test). Representative of 2 independent experiments.

**Fig. S4.**
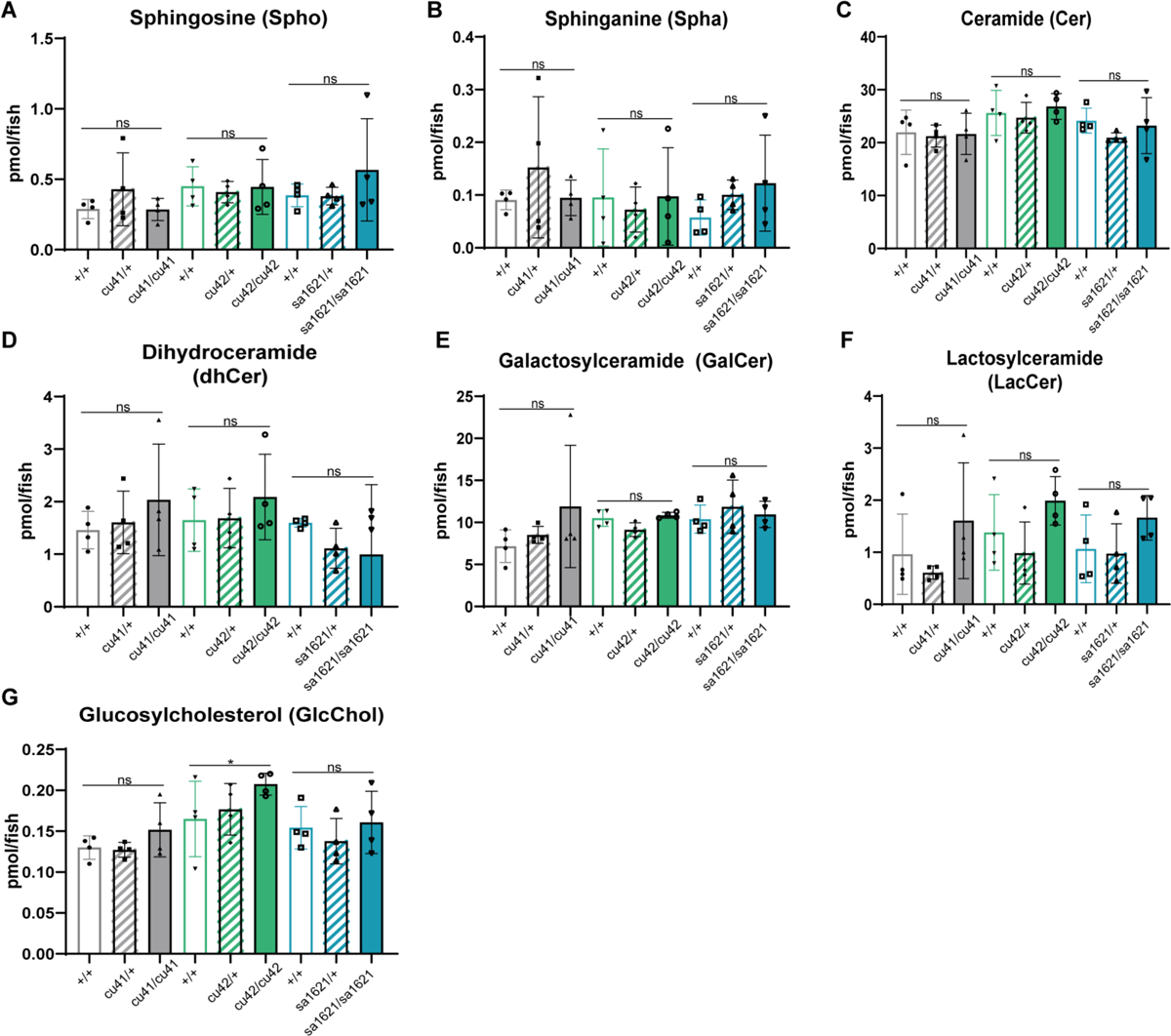
Quantification of lipids in 5dpf *gba1* mutants and their siblings.

**Fig. S5.**
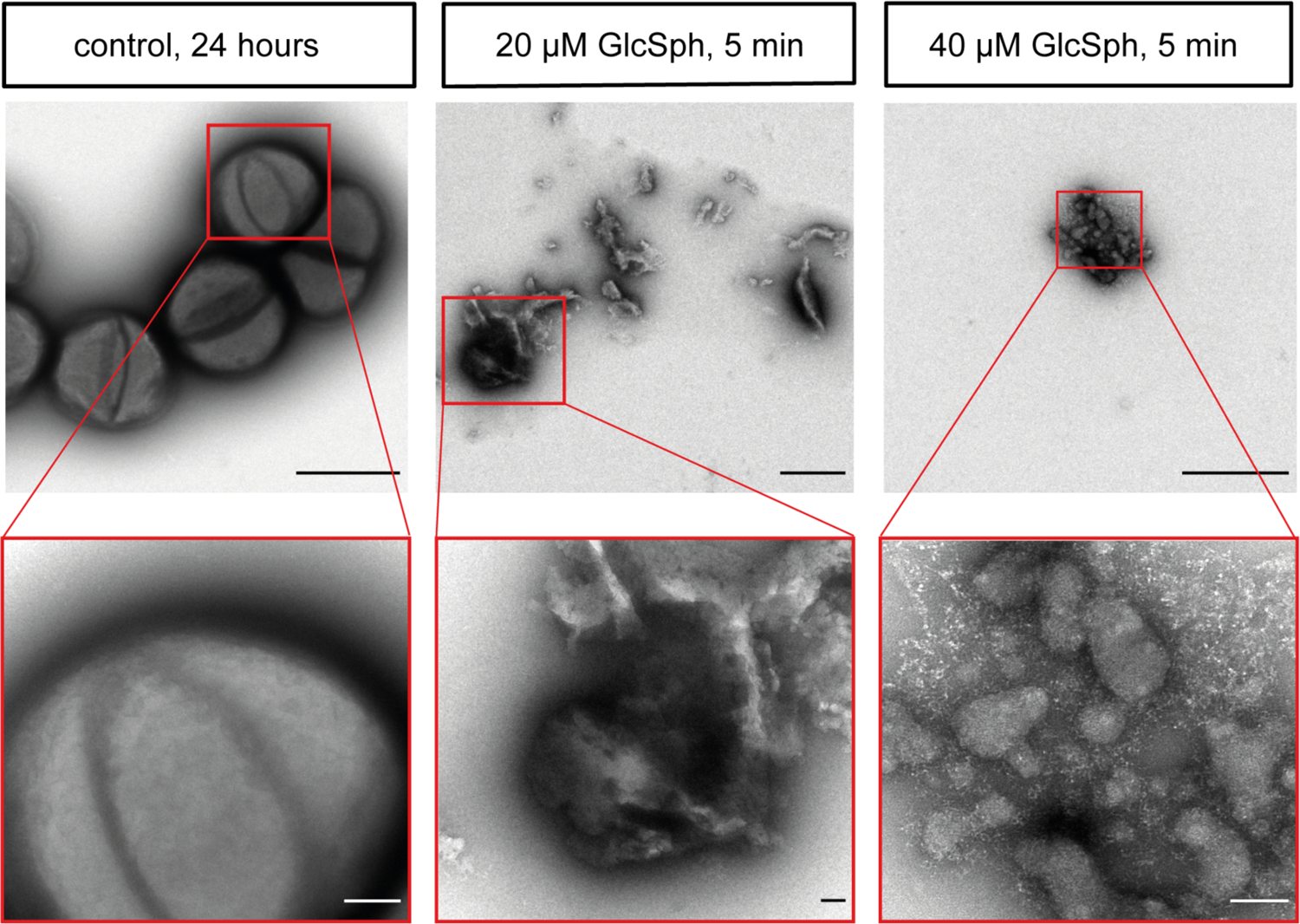
GlcSph-induced damage to *S. aureus* membranes. Representative negative stain-TEM images of *S. aureus* treated with GlcSph. Scale bar, top panel, 1 μm, bottom panel, 100 nm.

**Fig. S6.**
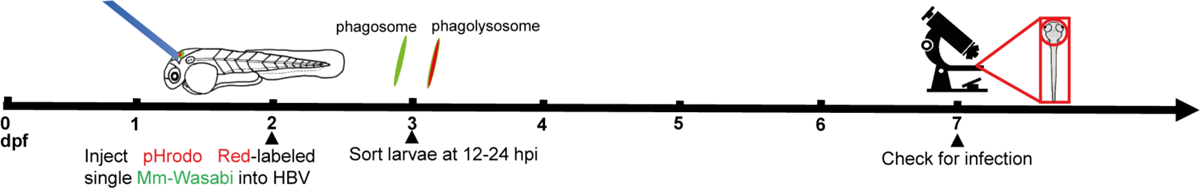
Schematic diagram of the infectivity experiment in the context of examining the effect of mycobacterial phagosomal versus phagolysosomal localization.

**Fig. S7.**
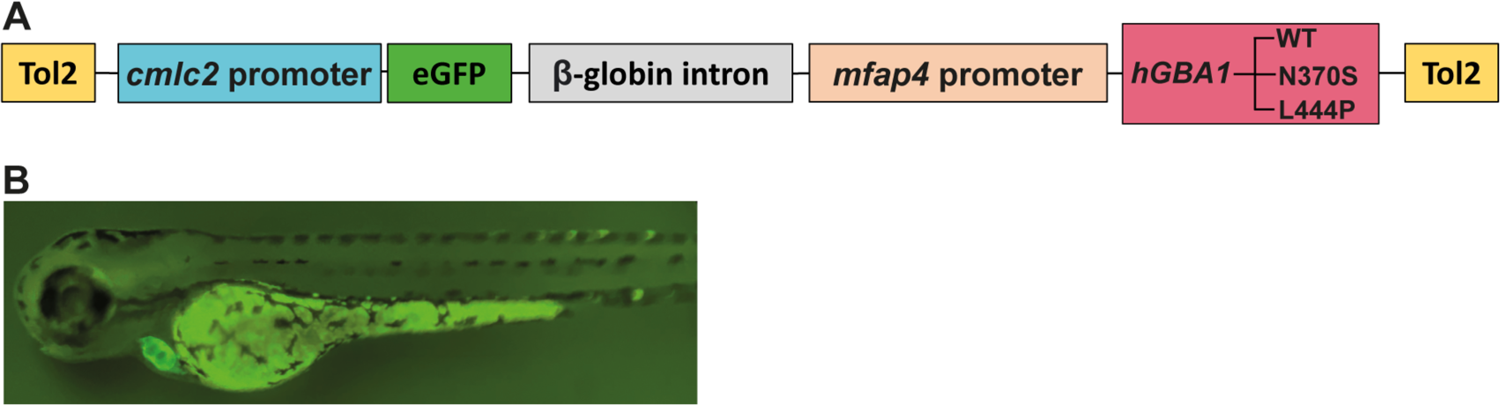
Transgenic zebrafish expressing human *GBA1* mutations in macrophages. *(A)* Schematic diagram showing the structure of the plasmid to generate the transgenic zebrafish line expressing macrophage-specific human *GBA1* mutations. *(B)* Representative image of the eGFP bleeding heart positive fish.

**Movie S1 (separate file).** Swimming abnormality (spinning) observed in 2.5 month-old *gba1^sa1621/sa1621^* mutant animals compared to normal swimming of their wild-type siblings. Of the four fish in the tank, the larger two are wild-type and the smaller two are *gba1^sa1621/sa1621^* mutants.

## Notes

https://www.dropbox.com/s/xq0t4uah6linw10/Supplemental%20Movie%201.mp4?dl=0

### Competing Interest Statement

The authors have declared no competing interest.

